# An Evolutionarily Conserved N-terminal Domain of RRF-3 Governs GTSF-1 Binding in Nematodes

**DOI:** 10.64898/2025.12.23.696163

**Authors:** Shamitha Govind, Sebastian Ruppert, Joseph Kirangwa, Virginia Busetto, Emily Nischwitz, Miguel Almeida, Svenja Hellmann, Hahn Witte, Ralf J. Sommer, Falk Butter, Sebastian Falk, Peter Sarkies, René F. Ketting

## Abstract

GTSF1 is an essential activating cofactor for PIWI proteins in many metazoans. In the nematode *Caenorhabditis elegans,* however, GTSF-1 does not bind PIWI, but associates with the RNA-dependent RNA polymerase RRF-3, supporting endo-siRNA (26G-RNA) biogenesis. Here, we demonstrate that this rewiring is deeply conserved across nematodes. For *C. briggsae* and *Pristionchus pacificus* we show that GTSF-1 interacts with RRF-3 and is essential for 26G-RNA production and fertility. We map this interaction to an N-terminal domain of RRF-3, termed the GTSF-1 interacting domain (GID), and show that the GTSF-1 zinc finger region alone is sufficient for binding. Mutagenesis identifies critical residues mediating this interaction and reveals that GTSF-1 stability depends on RRF-3. Other RdRPs possess GID-like domains, which we propose to bind GTSF-1-related proteins. Phylogenomic and structural analyses support GTSF-1–RRF-3 interactions across all major nematode lineages and map the shift in GTSF-1 activity to the last common nematode ancestor. We propose that binding of GTSF-1 induces conformational changes in RRF-3 that facilitate ERI complex assembly and activate RdRP function, paralleling its role as a PIWI activator.

## Introduction

RNA interference (RNAi) represents an ancient regulatory mechanism that employs antisense RNAs to modulate gene expression. Conserved across prokaryotes and eukaryotes, RNAi pathways serve dual roles in genome defense against viruses and transposons, and in developmental regulation of gene expression. In metazoans, RNAi pathways are classified into three distinct branches: the microRNA (miRNA), siRNA, and PIWI-interacting RNA (piRNA) pathways. RNAi machinery is comprised of three key enzymatic activities: RNA-dependent RNA polymerases (RdRPs) that synthesize double-stranded RNA (dsRNA), RNase III enzymes such as Dicer that process dsRNA into 20-30 nucleotide small interfering RNAs (siRNAs), and Argonaute/Piwi proteins, RNase H-like enzymes that utilize these small RNAs as sequence-specific guides to cleave or translationally repress target mRNAs.

Phylogenetic analyses suggest that the last common eukaryotic ancestor possessed all these three core RNAi components ^1,2^. However, gene duplications, losses, and functional innovations have diversified RNAi architecture across eukaryotic lineages. Plants and fungi, for instance, lack the PIWI-piRNA pathway but have evolved specialized RdRP-Dicer-Argonaute pathways that generate and amplify siRNAs^3^. Conversely, most metazoans have lost RdRPs, instead deriving their siRNAs from naturally occurring or exogenously introduced dsRNAs^4^, or from single-stranded precursors^5^. In metazoan lineages, Piwi-piRNA pathways have become specialized for transposon silencing in the germline^5^. However, certain parasitic flatworms, for example *S, mansoni* and *E. ocularis*, the dust mite *S. scabei,* and the majority of nematodes have independently lost the PIWI-piRNA pathway^6–8^. In contrast, nematodes and chelicerates (spiders, horseshoe crabs, scorpions) retain functional RdRP genes^7–11^. Despite these insights, our understanding of RNAi pathway evolution remains incomplete. While the core RNAi enzymes are well characterized, the auxiliary factors that modulate pathway specificity and efficiency are poorly understood, limiting our comprehension of how RNAi systems adapt and innovate across evolutionary time.

One such auxiliary factor is Gametocyte-specific factor 1 (GTSF1/Gtsf1), a conserved PIWI cofactor that was identified as essential for piRNA function and fertility in mice, flies, and silk moths^12–18^. GTSF1 proteins, which typically are between 20 and 30 kDa in mass, contain two tandem CHHC-type zinc fingers, each forming an independent fold around a central zinc ion-a structural motif also present in TRM13 tRNA modification enzymes and the minor spliceosomal protein U11-48K^19^. Absent from fungi and plants, GTSF1 proteins are deeply conserved across Metazoa, with many lineages encoding multiple paralogs^14,20–22^. Biochemical reconstitution experiments with Piwi proteins from mice, flies and silk moths have revealed that GTSF1 binds PIWI proteins engaged with target mRNA and induces conformational changes that generate catalytically active PIWI* complexes. For cytoplasmic Piwi proteins such as *M. musculus* MIWI and MILI or *B. mori* Siwi and Ago3, GTSF1 stabilizes a cleavage-competent conformation that facilitates efficient target mRNA slicing^14,16,23^. For catalytically inactive Piwi proteins, such as nuclear Piwi in *D. melanogaster*, GTSF1 binding creates protein-protein interaction surfaces that recruits chromatin-modifying complexes for transcriptional silencing ^24^.

The nematode *Caenorhabditis elegans* encodes a large complement of Argonaute proteins, including the canonical Piwi protein PRG-1^25–28^. PRG-1 binds piRNAs (termed 21U-RNAs in *C. elegans*), but unlike other metazoan Piwi proteins, it targets the entire germline transcriptome, including protein-coding genes and transposons ^25,26,29^. Loss of PRG-1 is not lethal but causes progressive fertility defects across generations^30^. Surprisingly, *C. elegans* GTSF-1 does not interact with PRG-1 or participate in the piRNA pathway. Instead, GTSF-1 associates with RRF-3, an RdRP that synthesizes 26G-RNAs, a distinct class of endogenous siRNA^31^. *C. elegans* GTSF-1 is required to assemble RRF-3 into a multiprotein complex named ERI, which includes Dicer, thereby coupling dsRNA synthesis with Dicer processing ^31,32^. These 26G-RNAs subsequently load onto the Argonautes ALG-3/4 and ERGO-1 to target protein-coding genes and pseudogenes^33,34^.

RRF-3 orthologs are widespread throughout the phylum Nematoda, whereas PRG-1 orthologs are often absent^8^. In basal nematodes, RRF-3-like RdRPs have been proposed to functionally replace the lost piRNA pathway by synthesizing siRNAs complementary to transposon sequences^8^. Indeed, nematode RNAi systems are heavily reliant on small RNA amplification by RdRPs. Unlike canonical metazoan RNAi systems, Argonaute-bound 21U-RNAs and 26G-RNAs in nematodes do not directly silence targets. Rather, they trigger the RdRPs RRF-1 and EGO-1 to synthesize abundant secondary siRNAs termed 22G-RNAs^30,35–38^. These 22G-RNAs then load onto nematode-specific WAGO Argonautes to induce transcriptional silencing and chromatin modification ^27,39,40^.

Within this intricate architecture of RNAi pathway, the molecular determinants of *C. elegans* GTSF-1 specificity for RRF-3 remain unclear. More fundamentally, the evolutionary origins and selective pressures that redirected *C. elegans* GTSF-1 from PIWI binding to RRF-3 binding are not understood. To address these questions, we performed a comparative analysis of GTSF-1 function in the nematodes *Caenorhabditis briggsae* and *Pristionchus pacificus*. Despite having diverged from *C. elegans* approximately 100 and 300 million years ago respectively, these species express 21U-, 22G-, and 26G-RNAs as well as the protein components of their respective pathways ^41–44^. These species provide an opportunity to probe the functional conservation of nematode GTSF-1 across deep evolutionary timescales.

In this work, we demonstrate that *C. briggsae* and *P. pacificus* GTSF-1 retain robust, specific association with RRF-3 and are essential for RRF-3-dependent 26G-RNA biogenesis. The zinc finger region of nematode GTSF-1 alone is sufficient for RRF-3 binding. GTSF-1 interacts with an N-terminal region of RRF-3, which we designate the GTSF-1 interacting domain (GID). AlphaFold modeling revealed that RRF-1 and EGO-1 also possess a GID-like fold/domain, which might be bound by GTSF-1-like proteins. We detected GTSF-1 and RRF-3 orthologs across the nematode phylum, and analysis of the predicted complex structures support a specific GTSF-1-RRF-3 interactions in multiple species but no interaction with PRG-1. Notably, structural predictions from a nematode sister lineage reveal a strong PIWI-GTSF-1 interaction, mapping the origin of RdRP binding to the nematode ancestor. We propose a conserved function for GTSF-1 to activate RRF-3 by inducing conformational rearrangements and promoting recruitment of the ERI complex. Finally, this work highlights a striking case of process homology within RNAi pathways, illustrating how RNAi components can diverge mechanistically while maintaining the overall regulatory outcomes of gene silencing and genome defense.

## Results

### Nematode GTSF-1 Is Required for Fertility and the 26G-RNA Pathway

Multiple GTSF1 paralogs are found across animal species^21^. For example, *D. melanogaster* encodes four (Asterix/Gtsf1, CG14036, CG32625, and CG34283)^12,13,22,24,45^, *M. musculus* has three (GTSF1, GTSF1-like, and GTSF2)^16–18^ and the silkmoth *B. mori* has two (Gtsf1 and Gtsf1-like) paralogs^14,15,46,47^. In contrast, *C. elegans* (Cel), *C. briggsae* (Cbr), and *P. pacificus* (Ppa) each possess a single ortholog of GTSF1. These nematode GTSF-1 orthologs retain the characteristic tandem CHHC zinc fingers in their N-terminal region and an approximately 100 amino-acid C-terminal tail that encompasses the ‘central’ region previously identified as essential for Piwi binding ^13,18^ (Figure 1A). AlphaFold3 (AF3) predicted structures of these orthologs closely match the solution structure of mouse GTSF1^45^ (Figure 1B and Extended Figure 1A). The tandem zinc fingers are arranged within two α-helices, connected by a flexible linker and the C-terminal region consists of α-helices interspersed by disordered stretches (Figure 1B and Extended Figure 1A). Additionally, PpaGTSF-1 contains an intrinsically disordered region (IDR) of approximately 100 amino acids at its N-terminus (Extended Figure 1A). As observed for metazoan GTSF1/Gtsf1 proteins ^21^, the zinc finger regions of nematode GTSF-1 are more conserved than the C-terminal region (Extended Figure 1B).

**Figure 1.**
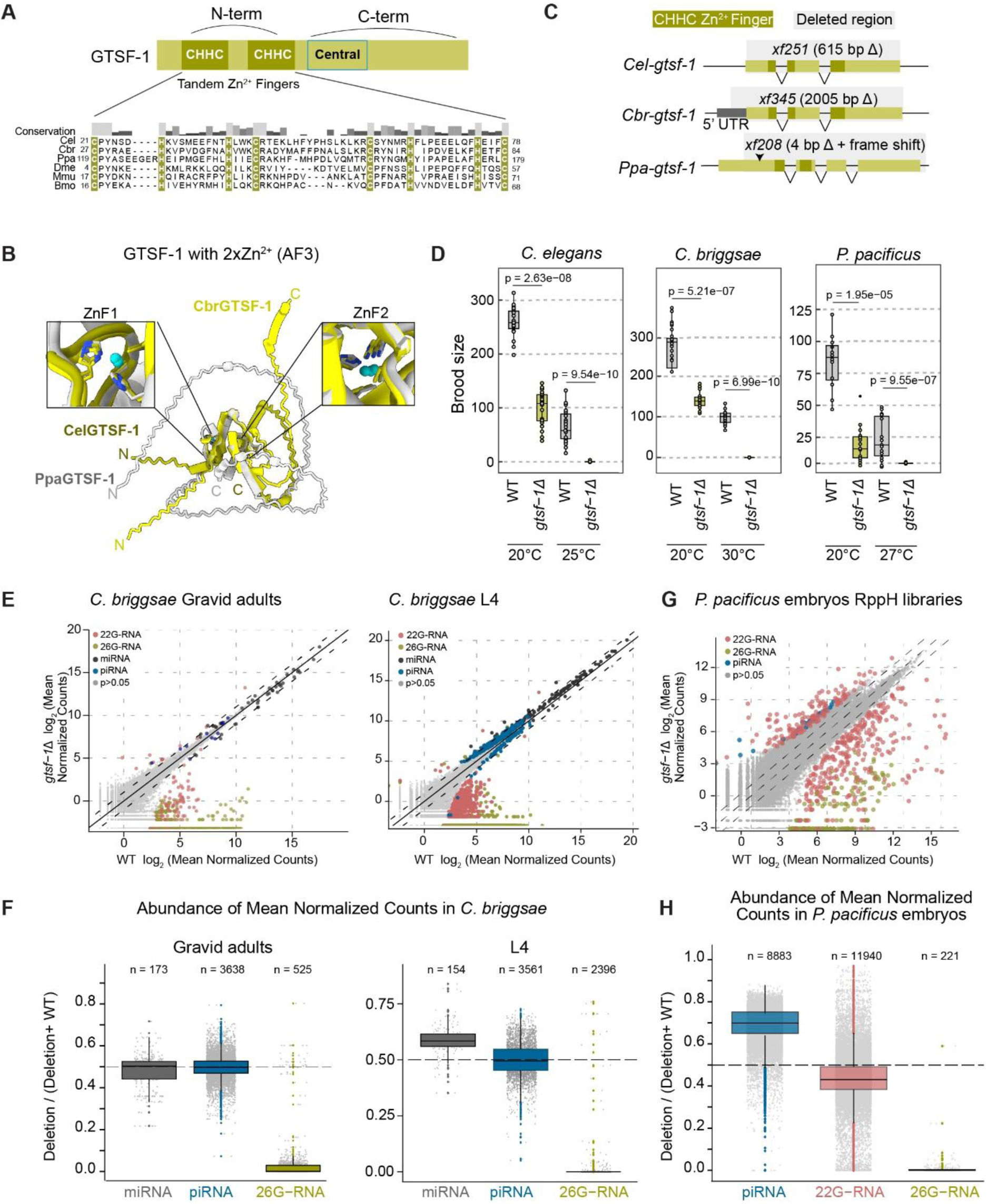
GTSF-1 is essential for fertility and 26G-RNA biogenesis (A) Domain architecture of GTSF-1 showing the N-terminal (N-term) zinc finger region, Central region and the C-terminal (C-term) region. Multiple sequence alignment (MSA) of residues spanning the CHHC zinc fingers in various GTSF-1 orthologs were performed using Clustal Omega and visualized with Jalview. The histogram above the alignment indicates conservation of physico-chemical properties in the MSA based on the AMAS method^127^. Species: *C. elegans* (Cel), *C. briggsae* (Cbr), *P. pacificus* (Ppa), *D. melanogaster* (Dme), *M. musculus* (Mmu), *B. mori* (Bmo). (B) AlphaFold 3monomer structure prediction of GTSF-1 bound to two Zn²⁺ ions. ZnF1 and ZnF2: zinc finger domains 1 and 2. Input sequences and prediction confidence scores are in Table 1. (C) Schematic of *gtsf-1* deletion alleles generated by CRISPR-Cas9. Green boxes indicate exons with darker shades showing genomic regions encoding the zinc fingers. Dotted lines mark deleted genomic regions. Arrow shows the position of a 4-bp deletion in *Ppa-gtsf-1* that results in frameshift and premature termination. (D) Brood size of wild-type (WT) and *gtsf-1* deletion animals at 20°C or 25-30°C (n = 15-25per genotype). P-values: Wilcoxon rank-sum test with Benjamini-Hochberg (BH) correction. (E, G) Small-RNA counts per gene in wild-type versus *gtsf-1* deletion animals for (E) *C. briggsae* gravid adult or L4animals and (G) *P. pacificus* embryos. Normalized counts (DESeq2) averaged across three biological replicates. Colored points: p &lt; 0.05.Solid line: equal expression; dotted lines: ±2 fold change. (F, H) Normalized count abundance per gene between wild-type and *gtsf-1* deletion samples by small-RNA type. (F) *C. briggsae* gravid adult or L4animals and (H) *P. pacificus* embryos. Dotted line: equal counts. n = number of genes per category.

In our previous work, we demonstrated that *C. elegans* GTSF-1 is essential for the 26G-RNA pathway ^31^. Mutants defective in this pathway exhibit impaired spermatogenesis, resulting in reduced brood size and sterility at elevated temperatures. To study this further, we generated *gtsf-1* mutant alleles in all three nematode species using CRISPR/Cas9 (Figure 1C). In *C. briggsae* and *C. elegans*, we created a complete gene deletion. In *P. pacificus*, our alleles preserve the first 100 amino acids in-frame before introducing a 4 base-pair deletion resulting in premature stop codon. The *gtsf-1* deletion allele in *C. elegans* showed reduced brood size at 20°C and sterility at 25°C (Figure 1D). We then tested whether *C. briggsae* and *P. pacificus gtsf-1* similarly contribute to fertility. In both species, *gtsf-1* deletion alleles reduced brood size at 20°C and caused complete sterility at stressful temperatures of 30°C and 27°C, respectively (Figure 1D).

In *C. elegans*, 26G-RNAs are expressed in the germline and loaded onto distinct Argonaute proteins depending on the gonadal context: ALG-3 and ALG-4 in spermatogenic gonads, or ERGO-1 in oogenic gonad and in embryos ^27,33,34,48^. We previously demonstrated that *gtsf-1* deletion in *C. elegans* eliminates both the ALG-3/4 and ERGO-1 classes of 26G-RNAs ^31^. To assess whether this function is conserved across nematodes, we sequenced small RNAs from *gtsf-1* mutants in *C. briggsae* and *P. pacificus*. As in *C. elegans*, *gtsf-1* mutants in *C. briggsae* displayed near complete loss of 26G-RNAs in both life stages, while piRNAs and miRNAs remained either unaffected or only marginally changed (Figure 1E and 1F). Additionally, 22G-RNAs downstream of 26G-RNA target genes were depleted, particularly in gravid adult stage animals (Extended Figure 1D and 1E). For *P. pacificus*, we analyzed RppH treated libraries from embryos of wild-type and *gtsf-1* deletion animals. The population of 26G-RNAs as well as the 22G-RNAs downstream of 26G-RNA target genes were strongly depleted upon loss of *gtsf-1* (Figure 1G and 1H; Extended Figure 1C). Interestingly, we also observed a moderate increase in piRNAs (Figure 1H).

Notably, a transgene expressing *C. briggsae* GTSF-1 in a *C. elegans gtsf-1* deletion strain partially rescued fertility defects, hypersensitivity to RNAi, and overall levels of 26G-RNAs and their downstream 22G-RNAs (Supplementary Figure 1A-D). Interestingly, the rescuing effect on brood-size was stronger at 20°C than 25°C. We conclude that the phenotypes we previously described for *gtsf-1* loss in *C. elegans* are conserved in *C. briggsae* and *P. pacificus*.

## Nematode GTSF-1 Associates with RRF-3

In *C. elegans*, the ERI Complex (ERIC) synthesizes 26G-RNAs through a coordinated multi-step process. The current hypothesis is that a NYN domain nuclease, RDE-8 or NYN-3, initiates the pathway by cleaving target RNAs upstream of RRF-3-mediated dsRNA synthesis^49,50^. RRF-3, assisted by the Tudor domain protein ERI-5 and the DExD/H-box helicase DRH-3, generates long double-stranded RNAs^32,48,51–56^. These dsRNAs are subsequently processed by Dicer, phosphatase PIR-1, and DEDDh-like exonuclease ERI-1 into 26-nucleotide RNAs with a 5’ monophosphate^57–59^. Additional factors, including RDE-4 and ERI-3, also participate in this process^32,60^.

Immunoprecipitation coupled with label-free quantitative mass spectrometry (IP-MS) on GTSF-1 in *C. elegans* adult extracts readily detects RRF-3 ^31^. IP-MS of epitope tagged GTSF-1 expressed via a transgene additionally enriches for ERI-5 and DRH-3^31^. To determine whether the GTSF-1-RRF-3 interaction is conserved across nematodes, we performed GTSF-1 IP-MS on adult extracts from *C. briggsae* and *P. pacificus*, using affinity-purified antibodies. *C. briggsae* GTSF-1 IP-MS significantly enriched RRF-3 and ERI-5 (Figure 2A). In *P. pacificus*, we performed two IP-MS experiments with different controls. Using anti-IgG antibodies as a negative control, GTSF-1 IP enriched RRF-3 and DRH-3, but not ERI-5 (Figure 2B: Left). This experiment also showed enrichment of the Argonaute ALG-1 (log₂FC = 1.89, log₁₀ p-value = 2.83), which is loaded with miRNAs in *C. elegans*^61^. To further validate antibody specificity, we compared GTSF-1 IPs from wild-type gravid adults to those from *gtsf-1* deletion animals. In this experiment, ERI-5 was now enriched in GTSF-1 IPs, in addition to RRF-3 and DRH-3 (Figure 2B: Right). Homologs of the 26G-RNA pathway components PIR-1 and two NYN domain proteins were also detected, although not enriched (Figure 2B: Left). However, ALG-1 was no longer detected, suggesting it is not a genuine GTSF-1-interacting protein.

**Figure 2.**
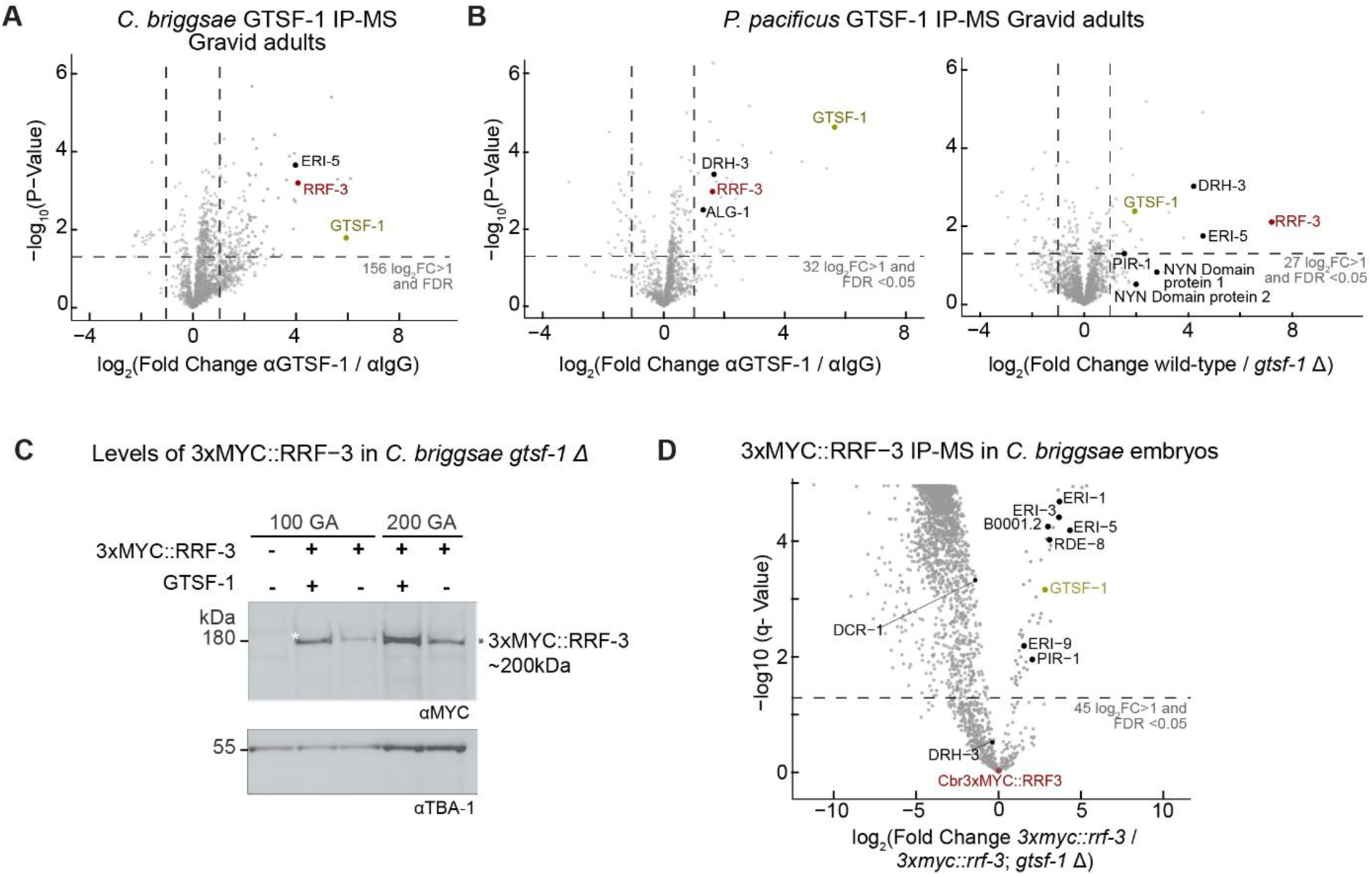
GTSF-1 interacts with RRF-3 in the ERI complex (A-B) Co-immunoprecipitation coupled to quantitative mass spectrometry (Co-IP/MS) of GTSF-1 on gravid adult animal lysates. Dotted lines: log₂ fold change ±1 and p &lt; 0.05. ERI complex and ALG-1 homologs highlighted. (A) *C. briggsae*; αCbrGTSF-1 antibodies with IgG control (B) *P. pacificus*; Right: αPpaGTSF-1 antibodies with IgG control, Left: αPpaGTSF-1 antibodies with IgG control. (C) Western blot of 3xMYC::RRF-3in *C. briggsae* wild-type or *gtsf-1* deletion animals (100 or 200 gravid adults (GA)). TBA-1: tubulin loading control. (D) Co-IP/MS of 3xMYC::RRF-3 using αMYC antibodies, on embryo lysates from *C. briggsae 3xmyc::rrf-3* versus *3xmyc::rrf-3; gtsf-1* deletion animals. X-axis: log₂ fold change; Y-axis: Adjusted p-values (q-value). Dotted lines: fold change (FC) ±2. ERI complex homologs highlighted.

In *C. elegans* GTSF-1 is required for coupling RRF-3 to ERIC. Immunoprecipitation of RRF-3 from embryo extracts enriches all known ERIC proteins in wild-type animals, but only ERI-5 in *gtsf-1* mutants ^31^. To test the conservation of this dependency, we generated epitope-tagged RRF-3 lines in *C. briggsae* and performed IPs in both wild-type and *gtsf-1* deletion backgrounds. Surprisingly, loss of *gtsf-1* caused a marked reduction in RRF-3 protein abundance (Figure 2C), which was not observed previously for *C. elegans* ^31^. To account for this, we normalized all protein abundances to RRF-3 levels when comparing genotypes. In RRF-3 IPs from wild-type *C. briggsae*, we detected several homologs of ERIC components-ERI-1, ERI-3, ERI-5, ERI-9, RDE-8, and PIR-1-whereas these proteins were absent in *gtsf-1* deletion samples (Figure 2D). However, DRH-3 and DCR-1 were present in both conditions and DCR-1 even showed mild enrichment in *gtsf-1* deletion samples, suggesting that *C. briggsae* GTSF-1 is dispensable for RRF-3 interactions with DRH-3 and DCR-1. Since this pattern was not observed in *C. elegans*^31^, these findings reveal species-specific aspects of ERIC assembly.

These results establish that GTSF-1 selectively associates with RRF-3 in both *C. briggsae* and *P. pacificus*, with no detectable interactions with other RdRPs (RRF-1, EGO-1) or Argonaute proteins including PRG-1. Further, the function of GTSF-1 in ERIC assembly remains conserved in *C. briggsae*, even though species-specific dependencies were detected.

## The C-terminal region of CelGTSF-1 Is Dispensable for Binding RRF-3 and the ERI Complex

Coimmunoprecipitation studies from mouse and flies have demonstrated that the central portion of Gtsf1 contains aromatic residues that are essential for Piwi binding ^13,18^. Although the Piwi binding aromatic residues are conserved in many animal GTSF1 proteins, nematode orthologs lack them entirely (Figure 3A). Additionally, a relatively conserved positively charged surface on the first zinc finger of mouse GTSF1 has been implicated in RNA binding and shown to be essential to activate PIWI^16,45^. This positively charged region is absent from our set of nematode species (Figure 3B), suggesting that RNA binding may be lost in nematode GTSF-1. This is consistent with previous report in which *in vitro* iCLIP on purified *C. elegans* GTSF-1 did not precipitate RNA beyond background levels^31^.

**Figure 3.**
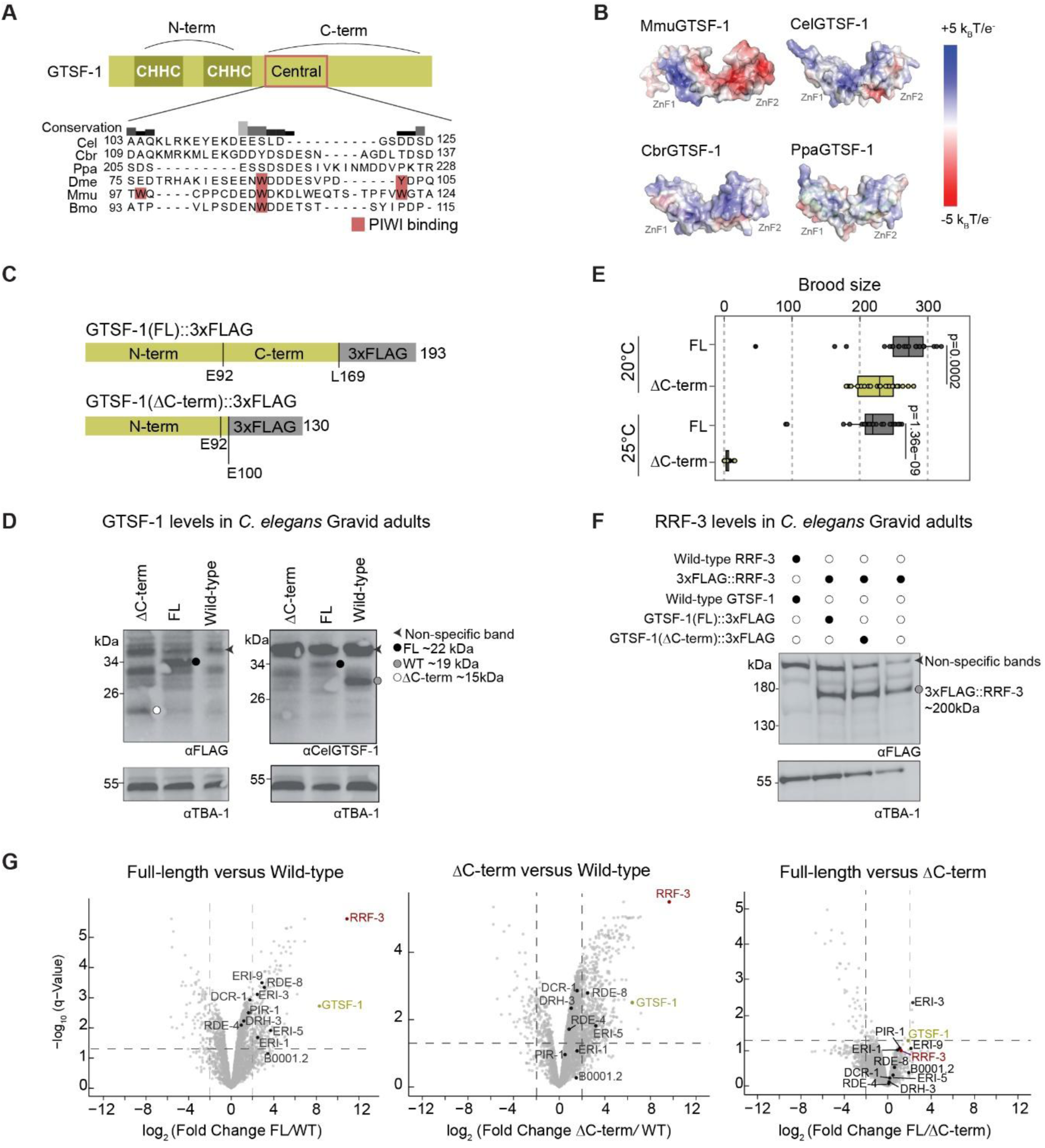
The C-terminal region of GTSF-1 is dispensable for RRF-3 interaction (A) Domain architecture of GTSF-1. MSA of Central region from various GTSF1/Gtsf1 orthologs highlighting aromatic residues implicated in PIWI binding. MSA performed with Clustal Omega; Visualization: Jalview; Species abbreviations and conservation histogram as in Figure 1A. (B) Electrostatic charge distribution on AF3 monomer GTSF-1 structures (-5kBT in red to +5kBT in blue). *M. musculus* GTSF1 exhibits positively charged ridge on ZnF1 as reported earlier^45^. *C. elegans* and *C. briggsae* display positive regions between ZnF1 and ZnF2, with reduced effect in *P. pacificus*. (C) Domain map of *C. elegans* GTSF-1mutants showing full-length (FL) or C-terminal truncation (ΔC-term) with 3×FLAG epitope tag positions. E92 and L169: end of N-terminal and C-terminal regions, respectively; E100: C-terminal truncation position. (D) Western blot detecting GTSF-1proteins with αFLAG and αCelGTSF-1 antibodies (100 gravid adults). αCelGTSF-1 targets C-terminal region of *C. elegans* GTSF-1, detecting the wild-type (WT) and, to some effect FL GTSF-1. Arrows: Non-specific bands. Filled circles: Positions of wild-type, FL or ΔC-term GTSF-1with expected sizes. TBA-1: tubulin loading control. (E) Brood sizes of GTSF-1mutant animals at 20°C and 25°C (n = 15-25). P-values: Wilcoxon rank-sum test with BH correction. (F) Western blot detecting 3×FLAG::RRF-3 in wild-type or GTSF-1 mutant animals (100 gravid adults). Filled or empty circles indicate presence or absence of indicated protein. Arrow: Non-specific band; Circle: 3×FLAG::RRF-3 with expected size. (G) Co-IP/MS for 3×FLAG::RRF-3 and GTSF-1::3×FLAG in *C. elegans* embryo lysates using αFLAG antibody. X-axis: log₂ fold change; Y-axis: Adjusted p-values (q-value). Genotype abbreviations: FL= *gtsf-1(fl)*::*3×flag*; *3×flag*::*rrf-3*; *rrf-3*(*pk1426*), ΔC-term= *gtsf-1*(*Δc*-*term*)::*3×flag*; *3×flag*::*rrf-3*; *rrf-3*(*pk1426*). Dotted lines: log₂ fold change ±2. ERI complex components highlighted.

We sought to determine how nematode GTSF-1 binds to RRF-3. First, we tested whether the C-terminal region of *C. elegans* GTSF-1 contributes to RRF-3 binding and ERI complex assembly. We generated a *gtsf-1* allele lacking this domain, leaving only the N-terminal region (N-term) fused to a 3×FLAG tag (Figure 3C). For brevity, we refer to this truncated GTSF-1 as ‘ΔC-term’. As a control, we also tagged full-length GTSF-1 with 3×FLAG. Western Blot analysis showed that wild-type GTSF-1 migrated slower than expected, as observed previously^31^ (Figure 3D). We detected the truncated GTSF-1 protein on a western blot, though at lower levels than full-length protein (Figure 3D). Interestingly, animals expressing GTSF-1 ΔC-term showed reduced fertility at 20°C (though they were more fertile than *gtsf-1* null mutants) and were near sterile at 25°C (Figure 3E). To test whether GTSF-1 truncation affects RRF-3 stability, we crossed these animals to a strain carrying a single-copy 3×*flag*::*rrf-3* transgene in a *rrf-3* deletion *(rrf-3*(*pk1426*)) background and assessed RRF-3 levels by Western blotting. The deletion of C-terminus did not affect RRF-3 protein levels (Figure 3F).

We next performed FLAG IPs on embryo extracts from animals carrying *3×flag::rrf-3; rrf-3(pk1426)* and expressing either full-length or truncated GTSF-1, also fused to 3×FLAG. In the presence of full-length GTSF-1, we enriched all known ERI complex proteins (Figure 3G), as expected. The enrichments of these proteins were significantly lower when GTSF-1 ΔC-term was expressed, but most of the factors were still detected (Figure 3G). These findings collectively demonstrate that the C-terminal regions of CelGTSF-1 are not essential for RRF-3 binding or ERI complex assembly, but they do appear to stabilize the assembled complex, especially at critical temperatures. This is consistent with previous report where recombinant GTSF-1 comprising only the N-terminal region was sufficient to precipitate RRF-3 from embryonic extracts^31^.

## Nematode GTSF-1 Binds a Conserved Folded Domain of RRF-3, the GID

Next, we aimed to identify the region of RRF-3 that interacts with GTSF-1. To achieve this, we generated AF3 Multimer models of GTSF-1 and RRF-3. For all three species, the models predicted that the CHHC-containing α-helices within the GTSF-1 N-terminal region contact a region in the N-terminus of RRF-3, distal to the RdRP catalytic site (Figure 4A and Extended Figure 2A; Average ipTM *C. elegans*= 0.636, *C. briggsae*= 0.646, *P. pacificus*= 0.686). This contrasts sharply with other metazoan GTSF1/Gtsf1 orthologs, which engage the catalytic PIWI domain of PIWI Argonautes ^23,24^. We refer to this region of RRF-3 as the GTSF-1-interacting domain or GID. We predicted the structures of isolated RRF-3 GID in complex with GTSF-1 and compared the results to full-length protein structures. Indeed, GTSF-1-GID complexes produced higher confidence scores than full-length GTSF-1-RRF-3 (Figure 4B). The structure of GID is conserved in *C. briggsae* and *P. pacificus* RRF-3 with 69%/30% identity and 79%/42% similarity to *C. elegans*, respectively (Figure 4C). GID conformation remained nearly identical between the full-length and isolated GID complexes (RMSD between RRF-3 FL and GID: *C. elegans*= 0.659 Å; RMSD *C. briggsae*= 0.718 Å; RMSD *P. pacificus*= 0.828 Å ) (Extended Figure 2B-2C).

**Figure 4.**
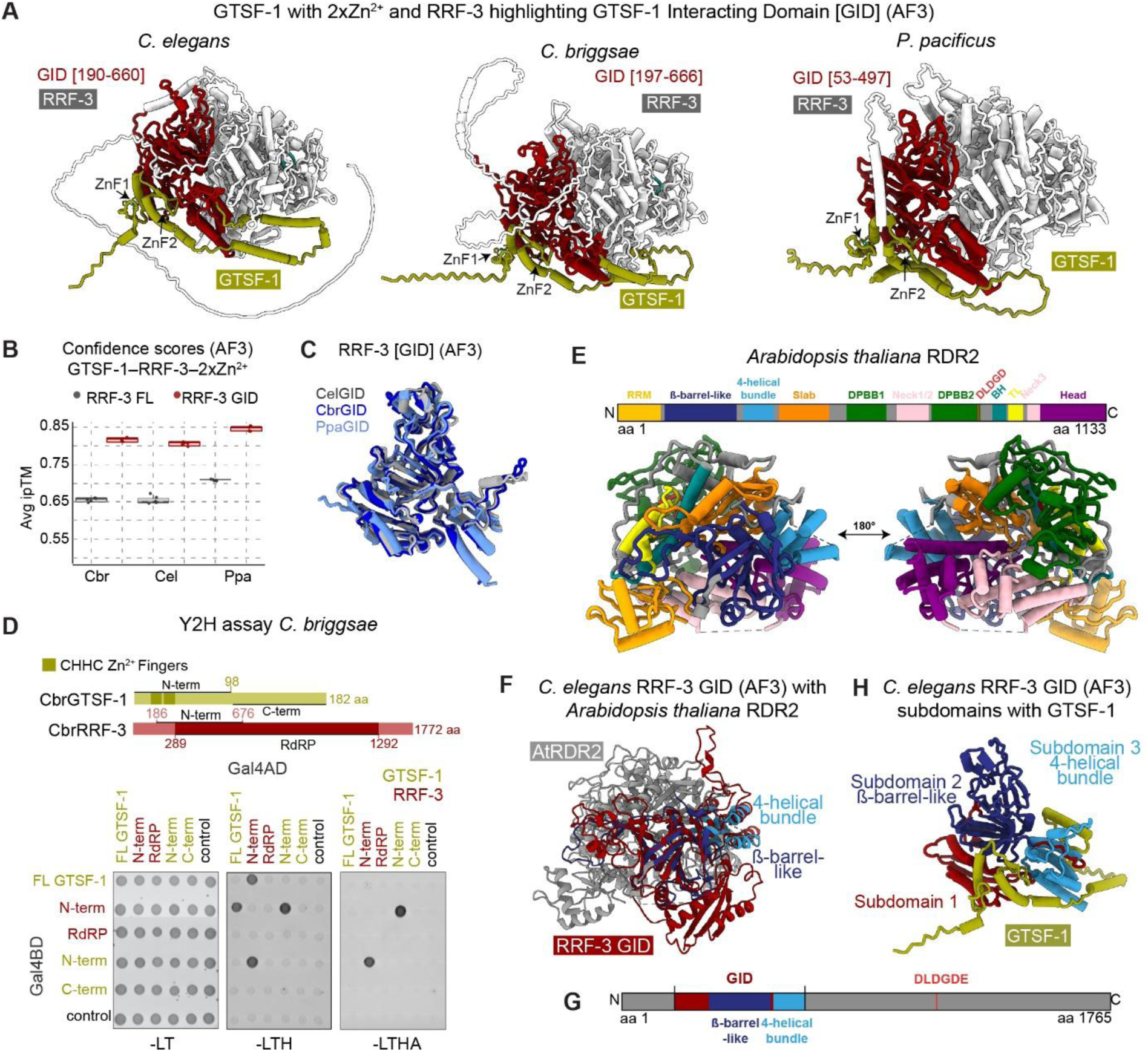
GTSF-1 N-terminus directly interacts with a conserved N-terminal region in RRF-3 (A) AlphaFold3multimer model of GTSF-1 (two Zn²⁺ ions) bound to RRF-3 for *C. elegans, C. briggsae* and *P. pacificus*. ZnF: zinc finger domains. GID: GTSF-1 Interacting Domain. Catalytic site (DLDGDE) highlighted in cyan. The respective GID is highlighted in red, with brackets indicating residue positions. (B) Average interface predicted template modeling scores (ipTM) from five AF3 models of RRF-3 full-length or GID paired with cognate GTSF-1 (two Zn²⁺ ions) orthologs (C) Superimposition of the RRF-3 GID from *C. elegans, C. briggsae* and *P. pacificus*, AF3 monomer. (D) Domain architecture of RRF-3 and GTSF-1 highlighting regions tested in Yeast Two-Hybrid assays. N-term and C-term: N-terminal and C-terminal regions. RdRP: InterPro RNA-dependent RNA polymerase domain. Yeast Two-Hybrid assays between GTSF-1 and RRF-3 regions under control (-LT: -Leu, -Trp), low stringency (-LTH: -Leu, -Trp, -His) and high stringency (-LTHA: -Leu, - Trp , -His, -Ade) selection. Control: Gal4 activation domain (AD) or DNA-binding domain (BD). (E) Crystal structure of *Arabidopsis thaliana* RNA-dependent RNA polymerase 2 (RDR2) from PDB (7RQS) with resolution of 3.57 Å. Domains and active site (DLDGD) are highlighted according to Fukudome et. al. 2021^63^. (F) *C. elegans* RRF-3 GID (AF3) superimposed on *Arabidopsis thaliana* RDR2 (AtRDR2). (G) Domain architecture of *C. elegans* RRF-3. The GID subdomains and RRF-3 active site are highlighted. (H) AF3 model of *C. elegans* RRF-3 GID in complex with GTSF-1 and 2x Zn²⁺, with its respective subdomains highlighted.

To validate the predicted structures, we tested pairwise interaction of regions of GTSF-1 and RRF-3 using the Yeast Two-Hybrid (Y2H) assay. For GTSF-1, we tested the full-length protein, the N-terminal region which contains both zinc fingers, and the C-terminal region (Figure 4D and Extended Figure 2D-E). The N-terminal region for PpaGTSF-1 also included the 100 amino acid IDR.

For RRF-3, we designed fragments spanning the GID at the N-terminal region and the InterPro/PFAM annotated RdRP domain. The annotated RdRP domain for *C. elegans* and *C. briggsae* RRF-3 partially overlapped with the region that was selected for the N-terminal region. We detected a robust interaction between GTSF-1 and the N-terminal region of RRF-3 for *C. briggsae*. This interaction was particularly strong for the N-terminal construct of GTSF-1, which showed positive interaction with RRF-3 even under stringent selection (Figure 4D). For *C. elegans* and *P. pacificus*, GTSF-1 constructs fused to Gal4 DNA-binding domain were self-activating (Extended Figure 2D-E). However, when expressed with the Gal4 activation domain, both full-length and N-terminal GTSF-1 showed positive interactions with the N-terminal region of RRF-3. For all three species, we did not detect any interaction with the RdRP construct. Altogether, our Y2H and structural analyses show that the GID is a conserved domain in the RRF-3 N-terminus that directly binds the zinc finger region of GTSF-1 in nematodes.

## GID Subdomains Share Homology with Eukaryotic RdRP N-Terminal Domains

To place RRF-3 GID in broader evolutionary context, we compared AF3-predicted RRF-3 structures to experimentally determined structures of plant RdRPs from *Arabidopsis* (AtRDR2)^62,63^ (Extended Figure 2F). The AtRDR2 structure comprises an N-terminal region containing an RRM-like fold, a β-barrel–like domain, and a four-helical bundle domain, followed by an active site cleft composed of multiple conserved domains including the slab, DPBB1 and DPBB2 (double-ψ β-barrel) domains, connector helices (neck 1, 2, and 3), bridge helix, trigger loop, and head domain^62,63^ (Figure 4E). Structural alignment revealed that the RRF-3 GID superimposes well with AtRDR2 and can be divided into three subdomains (Figure 4F-H). Subdomains 2 and 3 are structurally homologous to the N-terminal regions of AtRDR2 that precede the DPBB core. Subdomain 2 corresponds to the β-barrel-like fold, also referred to as the "PH" domain in ZmRDR2^64^ or the "Guide" domain in an older annotation of AtRDR2^62^. Subdomain 3 aligns with a four-helix bundle that is annotated as part of the "Slab" domain in both plant RdRPs (Figure 4G-H). In contrast, subdomain 1 of the GID did not show clear structural homology to plant RdRP domains, suggesting it may be nematode-specific (Figure 4H). Thus, RRF-3 GID is comprised of deeply conserved RdRP structural elements.

## Critical Residues Mediating the GTSF-1-RRF-3 Interaction

We next investigated the specific residues that comprise the GTSF-1-RRF-3 binding interface. Using the AF3 predicted structures, we identified key aromatic (W39) and positively charged amino-acids (R42, R63) adjacent to the zinc fingers on GTSF-1 that bind to RRF-3 GID (Figure 5A). ConSurf analysis using an alignment of several *Caenorhabditis* GTSF-1 sequences revealed good conservation of these residues (Figure 5A and Extended Figure 3A). Using CRISPR/Cas9, we created point-mutations in *C. elegans* GTSF-1 generating two strains carrying either W39A;R42A or R63A alone and assessed their fertility. GTSF-1 W39A;R42A mutants produced reduced brood size at 20°C but became sterile at 25°C (Figure 5B). In contrast, the brood-size of R63A mutants was comparable to wild-type animals at both temperatures (Figure 5B). However, we could not detect W39A;R42A GTSF-1 protein by Western blot, whereas R63A maintained wild-type protein levels (Figure 5C). Mutating W39 and R42 thus appear to destabilize GTSF-1. Given the close physical association between GTSF-1 and RRF-3, we reasoned that the observed degradation of GTSF-1 might be due to an inability to bind RRF-3. Consistent with this hypothesis, we did not detect wild-type GTSF-1 in RRF-3 deletion animals (Figure 5C).

**Figure 5.**
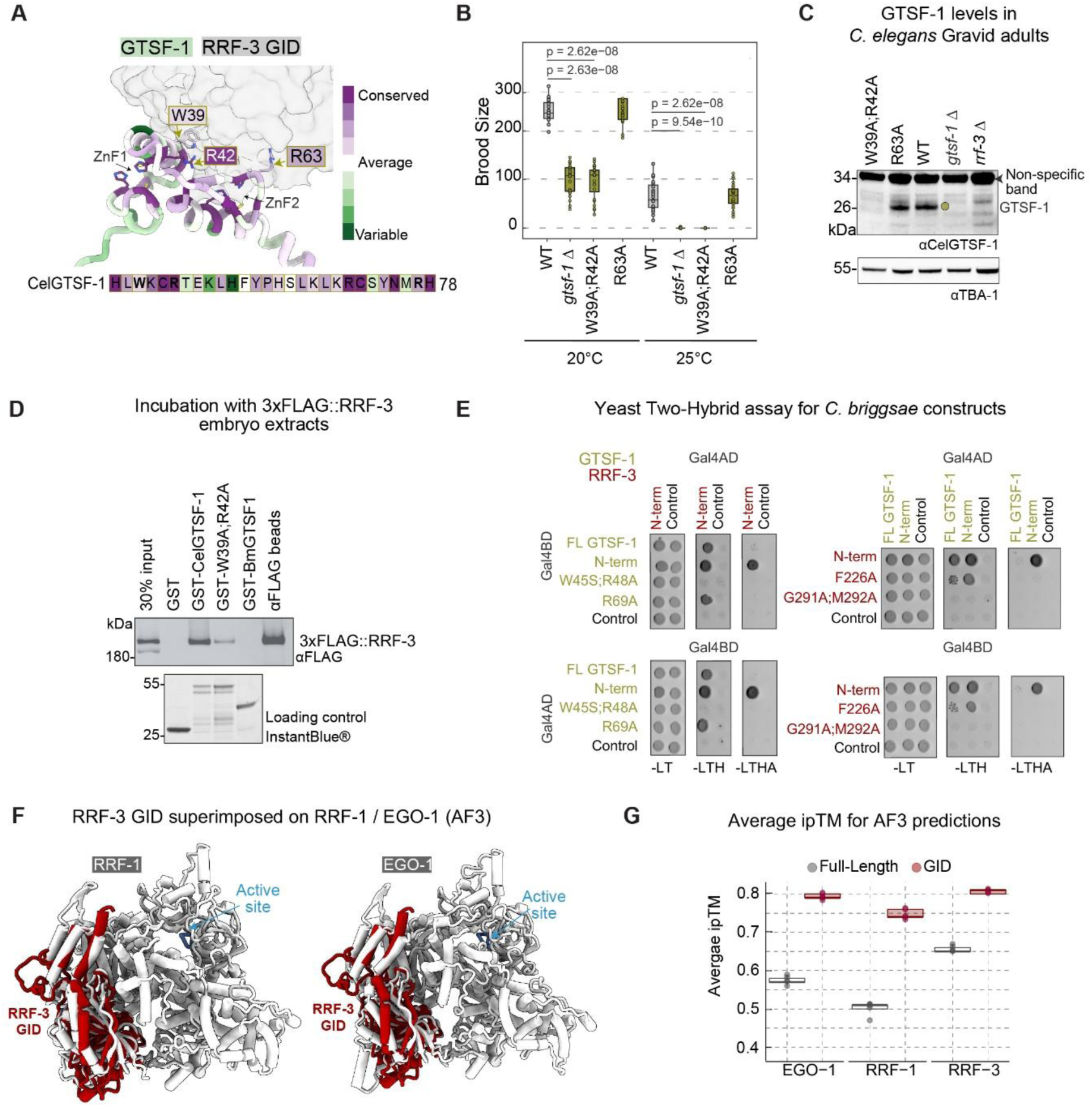
GTSF-1-RRF-3 interface residues are critical for interaction and fertility (A) *C. elegans* GTSF-1-RRF-3-2×Zn²⁺ complex (AF3) zoomed into the interaction interface. Residues on GTSF-1 predicted to bind RRF-3 GID displayed in stick format colored by ConSurf conservation score calculated using an MSA of 26 nematode species. Details in Table 2. (B) Brood size of *gtsf-1* point mutants, (n = 15-25). P-values: Wilcoxon rank-sum test with BH correction. (C) Western blot on GTSF-1 mutant animals compared to Wild-type (WT), *gtsf-1* deletion *(xf251)* and *rrf-3* deletion (*pk1426*) animals (100gravid adults). TBA-1: tubulin loading control. (D) Western blot displaying levels of 3×FLAG::RRF-3precipitated from *C. elegans* embryo lysates using GST or GST-GTSF-1 proteins. Input: 30% of embryo lysate used per pulldown. αFLAG beads: Positive control. Loading control: 10% of GST or GST-GTSF-1 proteins used for pulldown, visualized using InstantBlue®. (E) Yeast Two-Hybrid assays between *C. briggsae* GTSF-1 and RRF-3 with point mutations in the N-terminal constructs of both proteins. Constructs assayed under control (-LT: -Leu, -Trp), low stringency (-LTH: -Leu, -Trp, -His) and high stringency (-LTHA: -Leu, -Trp, -His, -Ade) selection. Control: Gal4AD or Gal4BD empty vectors serving as negative controls. (F) AF3Monomer structures of RRF-1 and EGO-1 superimposed with RRF-3. For RRF-3, only the GID region is visualized. Active site: DLDGE catalytic residues of RdRPs. RMSD between RRF-3 GID and RRF-1 = 0.980 Å over 139 pruned atom pairs (across all 305 pairs = 5.302) and EGO-1 = 0.944 Å over 138 pruned atom pairs (across all 308 pairs = 9.336). (G) Average ipTM scores from five AF3 models of GTSF-1 (two Zn²⁺ ions) with different full-length RdRPs and their respective GID.

We also performed GST pulldown assays using recombinantly expressed GST-fused GTSF-1 proteins. Both wild-type and W39A;R42A GTSF-1 were expressed at comparable levels in *E. coli*, indicating that the mutations do not intrinsically affect protein stability (Figure 5D, Bottom). However, when we incubated these proteins with *C. elegans* embryo lysates, wild-type GTSF-1 bound to 3×FLAG::RRF-3 much stronger than GTSF-1(W39A;R42A) (Figure 5D, Top).

Finally, we tested the effect of corresponding mutations on the CbrGTSF-1-RRF-3 interaction in a Y2H assay (Figure 5E and Extended Figure 3B). Consistent with the *C. elegans* data, CbrGTSF-1(W45S;R48A) did not bind to CbrRRF-3, while the CbrGTSF-1(R69A) did (Figure 5E). Through the AF3 predicted structures, we identified residues in the *C. briggsae* RRF-3 GID (F226, G291, and M292) that bind to W45 and R48 (Extended Figure 3B-C). Among these, G291 and M292 showed greater conservation than F226. When we tested these residues in Y2H, the G291A/M292A double mutation abolished the interaction between RRF-3 and GTSF-1 (Figure 5E). F226A also showed a weak but noticeable binding defect with GTSF-1. We conclude that the interface between GTSF-1 and RRF-3 that is predicted by AF3 is indeed required for their interaction, and that nematode GTSF-1 stability depends on its interaction with RRF-3.

## RRF-1 and EGO-1 possess GID-like domains that are predicted to interact with GTSF-1

*C. elegans* also encodes the RdRPs RRF-1 and EGO-1, which act downstream of RRF-3 but function independently of DCR-1 and the ERI Complex. Having identified a GTSF-interacting domain (GID) in RRF-3, we next asked whether RRF-1 and EGO-1 contain a similar structural feature. AlphaFold3 (AF3) models revealed that the N-terminal regions of RRF-1 and EGO-1 superimpose well with the RRF-3 GID (Figure 5F), suggesting a conserved fold across the three RdRPs. When AF3 was used to predict full-length complexes of GTSF-1 with each RdRP, the resulting confidence scores were lower for RRF-1 and EGO-1 than for RRF-3 (Figure 5G). Nevertheless, GTSF-1 was consistently positioned at the putative GID region for RRF-1 and EGO-1 (Extended Figure 3D–E). Notably, restricting predictions to only the GID regions markedly improved confidence (Figure 5G), and the resulting RRF-1 and EGO-1 GID–GTSF-1 complexes closely resembled the RRF-3 GID–GTSF-1 interface (Extended Figure 3F). We therefore propose that all three *C. elegans* RdRPs share a GID-like N-terminal architecture capable of interaction with GTSF-1-like proteins. The *in vivo* specificity of GTSF-1 for RRF-3 likely arises from additional factors that we can currently not identify.

## Conservation Patterns of GTSF-1, PIWI, and RdRP Across Phylum Nematoda Reveal Clade-Specific Losses

Having established the molecular basis for GTSF-1’s selectivity for RRF-3, we next sought to trace the evolutionary history of this interaction and determine whether nematode GTSF-1 originally may have partnered with PIWI or RdRP proteins. The phylum Nematoda comprises three classes: Dorylaimia (Clade I), Enoplia (Clade II), and Chromadoria (Clade C)^65^. Within Chromadoria, the order Rhabditida includes three subclasses: Spirurina (Clade III), Tylenchina (Clade IV), and Rhabditina (Clade V)^65^. Clade V contains the genera *Caenorhabditis and Pristionchus*. We searched for orthologs of *C. elegans* PRG-1, RRF-3, RRF-1/EGO-1, and GTSF-1 in species representing all major nematode clades. Our set included species from Enoplia and early-branching Chromadorea. We also included four species from phylum Nematomorpha, as they represent the sister group to Nematoda.

Through a reciprocal BLAST approach, we identified orthologs of PIWI, RdRP, and GTSF-1 in genomes of 80 nematode species (Figure 6A). Our results were consistent with previous reports: *C. elegans* PRG-1 orthologs were absent in Clades III and IV, as well as in several Clade I and II species, indicating repeated loss of the piRNA pathway across the nematode phylum^8^. RRF-3 like RdRPs are found across all clades, while RRF-1/EGO-1 proteins are mostly found within Clades III-V. Notably, we found both RRF-1 and EGO-1 only in *C. briggsae*, *C. brenneri,* and *C. elegans*, which suggests these gene paralogs likely arose after a recent gene duplication in *Caenorhabditis*. These results indicate that RRF-1/EGO-1 most likely lost the ability to bind GTSF-1, rather than that RRF-3 uniquely gained the property to bind GTSF-1.

**Figure 6.**
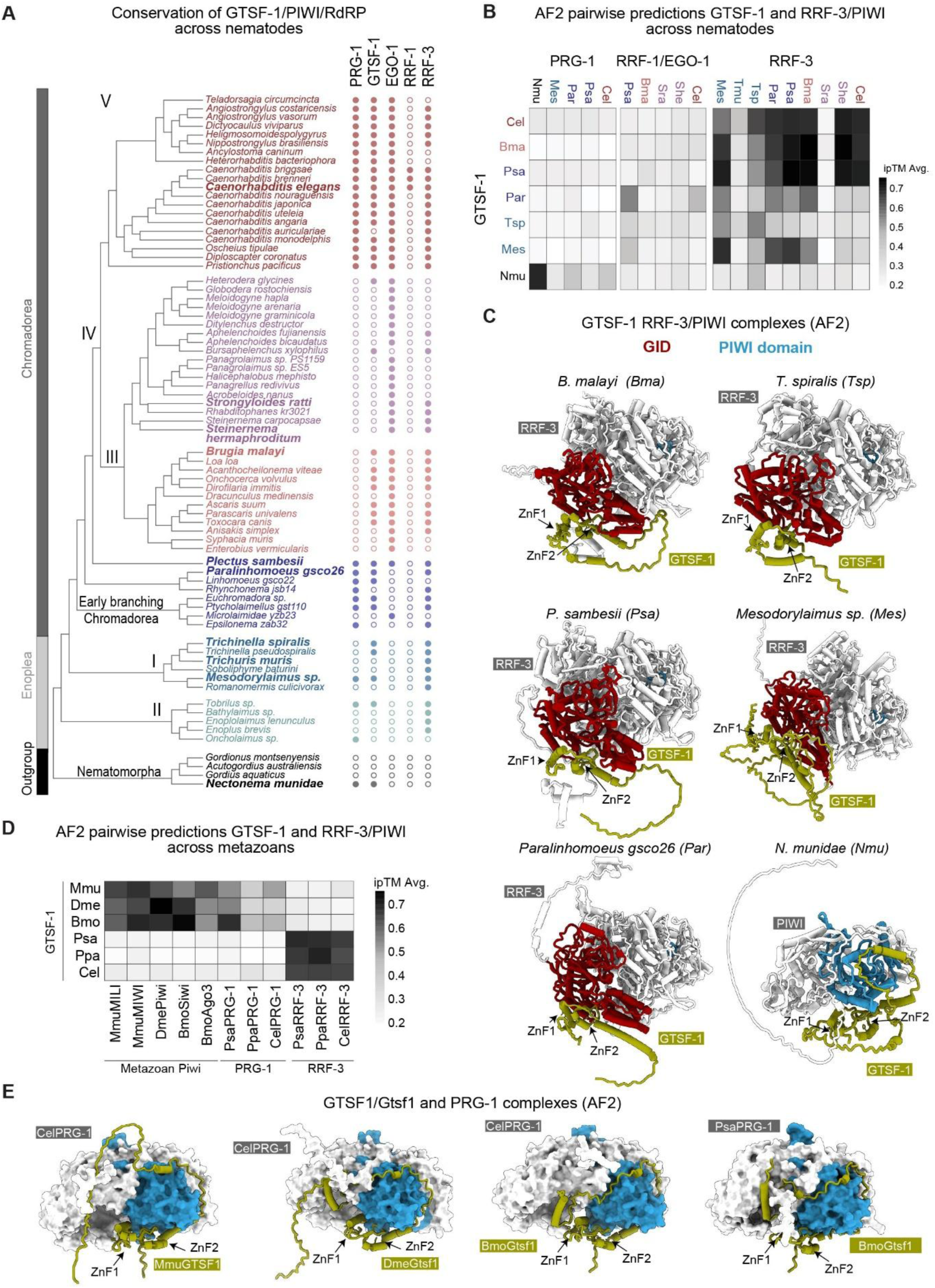
GTSF-1-RRF-3 interaction is conserved across nematodes (A) Conservation of selected proteins across phylum Nematoda and Nematomorpha. Species are color coded by Clades I-V and Early-branching Chromadorea. Filled and empty circles indicates presence or absence of indicated orthologous protein. Bold faced species were selected for AlphaFold2 Multimer pairwise prediction screen. (B) Heatmap of AlphaFold2 Multimer pairwise interactions between PIWI, RRF-1/EGO-1, and RRF-3 orthologs and GTSF-1 proteins. Species: She, *S. hermaphroditum*; Sra, *S. ratti;* Bma, *B. malayi*; Psa, *P. sambesi*; Par, *Paralinhomoeus gsco26*; Tsp, *T. spiralis*; Tmu, *T. muris*; Mes, *Mesodorylaimus sp*.; Nmu, *N. munidae*. (C) AF2 structures of cognate RRF-3-GTSF-1 or PIWI-GTSF-1 complexes from panel B. (D) Heatmap of AF2Multimer pairwise interactions between PIWI, PRG-1 and RRF-3 orthologs and GTSF-1/GTSF1/Gtsf1 proteins. Species additional to panel B: Mmu: *M*. *musculus*, Dme: *D. melanogaster*, Bmo: *B.mori* (E) AF2 structures of cognate PRG-1-GTSF1/Gtsf1 complexes from panel D.

GTSF-1 showed patchy conservation across the phylogeny, though we detected it in at least one species from each clade (Figure 6A). Conservation was strongest within Clade V, where we found GTSF-1 in 19 of 20 species tested. In contrast, Clade IV showed the most striking loss: GTSF-1 was absent from 16 of 18 species, 11 of which also lacked RRF-3. Within the entire set, we identified 11 species where RRF-3 was present without GTSF-1. We also found 5 species where GTSF-1 was detected while RRF-3 was not. These phylogenetic patterns show that GTSF-1 and RRF-3 orthologs are present broadly across Nematoda, where they are found in most species, and that they tend to co-exist. However, gene loss seems to be frequent, indicating fast-changing small RNA pathway dynamics amongst nematodes. Finally, our set of Nematomorphs was largely devoid of PIWI, GTSF-1 and RdRPs. However, in one species - *Nectonema munidae* we did find an ortholog of both PIWI and GTSF-1.

## Structural Modeling Reveals Specificity of GTSF-1 Interactions

We extracted high-confidence sequences from key species and used AlphaFold2 Multimer to predict pairwise interactions between GTSF-1 proteins and either RdRPs or PIWI proteins. We selected 10 species representing key phylogenetic positions (Figure 6A: Bold-face). *Trichinella spiralis* (Clade I) and *Brugia malayi* (Clade III) both contained GTSF-1 and RRF-3 but lacked PRG-1. Two Clade IV species (*Strongyloides ratti* and *Steinernema hermaphroditum*) and the Clade I species *Trichuris muris* encoded RRF-3 but no GTSF-1. Three species were of particular interest: the Clade I species *Mesodorylaimus sp.* and the early-branching Chromadorea species *Paralinhomoeus gsco26 and Plectus sambesi.* All three encoded PRG-1, RRF-3, and GTSF-1, allowing us to test both potential interactions. Finally, *N. munidae* GTSF-1 and PRG-1 sequences enabled us to predict GTSF-1’s binding partner in the phylum most closely related to the Nematoda.

The GTSF-1 sequences varied considerably in length-*T. spiralis* GTSF-1 comprised only 95 amino acids restricted to the zinc finger region, whereas *Mesodorylaimus* GTSF-1 extended to 263 amino acids with a long C-terminal (Extended figure 4A). Despite this variation, all sequences retained the tandem zinc fingers arranged within α-helices. Interestingly, *T. spiralis*, *Mesodorylaimus sp.,* and *Paralinhomoeus* GTSF-1 proteins contained CCHC motifs instead of the canonical CHHC motif, and AF3 predicted these to fold into functional zinc fingers (Extended Figure 4B).

We detected high-confidence interactions between cognate GTSF-1 and RRF-3 proteins from *B. malayi, P. sambesii*, and *Mesodorylaimus sp*., with lower but still favorable scores for *T. spiralis* and *Paralinhomoeus* (Figure 6B). To test the specificity of these predicted interactions, we compared closely related RRF-3 orthologs from *T. spiralis*, which does have a GTSF-1 homolog, and *T. muris*, which does not encode GTSF-1. *T. spiralis* RRF-3 was predicted to bind GTSF-1 homologs from various nematodes, also from other clades, such as *C. elegans*. In contrast, *T. muris* RRF-3 showed no favorable binding with any of the GTSF1 proteins from our dataset, also not from the closely related *T. muris*. This result demonstrated that AlphaFold2 homology modeling predicted differences in interaction specificity even among closely related orthologs. All favorable GTSF-1-RRF-3 complexes featured a GID in the N-terminus of RRF-3 that engaged the GTSF-1 NTD, mirroring the architecture we observed in *C. elegans* (Figure 6C).

Within Clade IV, where most species lack GTSF-1, the two species we tested displayed opposite patterns. *S. hermaphroditum* RRF-3 achieved surprisingly high scores with several GTSF-1 proteins, even if the species itself does not have GTSF-1 anymore. In contrast, *S. ratti* RRF-3 scored very low across all tested combinations. Closer inspection revealed that *S. ratti* RRF-3 lacked portions of the GID (Extended Figure 4D). This further underscores the critical role of this domain in mediating GTSF-1 binding. The high GTSF-1 binding propensity of *S. hermaphroditum* RRF-3 suggests that GTSF-1 was either recently lost or that a distant GTSF-1 homolog, which we failed to identify, still binds the RRF-3 GID.

RRF-1/EGO-1 like proteins generally did not show reliable interaction with GTSF-1 proteins. Even in *C. elegans*, the predicted interaction between GTSF-1 and RRF-1/EGO-1 with AF2 showed substantially lower confidence scores compared to the RRF-3–GTSF-1-2xZn^2+^ complexes in AF3 predictions. However, there were a few exceptions. The most notable was PsaRRF-1 with ParGTSF-1, which does not appear to be an artifact, as ParGTSF-1 binds PsaRRF-1 at the correct location-the GID (Extended Figure 4E). Even though this is clearly not biologically significant, and since PsaGTSF-1 itself does not bind PsaRRF-1, this finding again hints that RRF-1 proteins may be capable of accommodating GTSF-1 like proteins. Finally, none of the PRG-1 proteins showed reliable interaction predictions with any of the nematode GTSF-1 proteins. In contrast, PRG-1 from *N. munidae*, from the Nematomorpha phylum, exhibited a high-confidence interaction with NmuGTSF-1.

These findings demonstrate that nematode GTSF-1 maintains a conserved structural interface with RRF-3 GID, even over substantial evolutionary distances. We found reliable GTSF-1-RRF-3 predictions in at least one species from each major nematode lineage, including Enoplea, the sister group to all other nematodes. Notably, even in species encoding both PIWI and RRF-3, GTSF-1 preferentially interacted with RRF-3. In contrast, Nematomorpha-the sister group to Nematoda-did not encode RdRPs and showed strong GTSF-1-PIWI predictions. Thus, the functional divergence of GTSF-1 to bind RRF-3 and not Piwi is specific to Nematoda.

## Differences in nematode GTSF-1 responsible for loss of PIWI binding

We show that GTSF-1-RRF-3 interaction is preferred over GTSF-1-PIWI throughout nematodes. Based on these conservation patterns, the most parsimonious explanation is that GTSF-1-RRF-3 interaction was already present at the base of the nematode phylogeny. Interestingly, despite PIWI loss in numerous lineages, several nematode species have retained this Argonaute. This raises the question: did nematode PIWI proteins diverge, preventing GTSF-1 binding, or is nematode GTSF-1 incompatible with PIWI recognition?

To address this, we tested pairwise interactions between GTSF-1 and either RRF-3 or PIWI proteins from *B. mori*, *D. melanogaster*, *M. musculus* and the nematodes *C. elegans*, *P. pacificus* and *P. sambesi*. As expected, nematode GTSF-1 showed high confidence interaction scores with RRF-3 orthologs but low scores with all PIWI proteins, including PRG-1 orthologs (Figure 6D). In contrast, non-nematode Gtsf1 proteins showed favorable interaction scores with all PIWI proteins, including nematode PRG-1, but very low scores with nematode RRF-3 proteins (Figure 6D). The lower ipTM scores with PRG-1 could be due to differences in C-terminal regions of GTSF-1/Gtsf-1 proteins, as this region has been shown to mediate specificity to cognate Piwi proteins, even among paralogs^16^. Overall, we find these cross-phylum predictions reliable because the predicted Gtsf1-PRG-1 complexes closely resemble Gtsf1-PIWI structures from other animals^23,24^, in that GTSF1/Gtsf1 binds PRG-1 at the Piwi domain with the central and C-terminal regions wrapped around it (Figure 6E and Extended Figure 4F).

These reciprocal interaction patterns reveal that nematode PRG-1 retains the structural capacity to bind non-nematode Gtsf1 proteins, while non-nematode Gtsf1 proteins cannot recognize nematode RRF-3. This strongly suggests that nematode PIWI proteins did not lose their GTSF1-binding potential. Instead, nematode GTSF-1 underwent evolutionary changes that eliminated its PIWI-binding capacity while establishing a specific interaction with RRF-3. This functional divergence of GTSF-1 identified within the phylum Nematoda shows a remarkable example of protein partnership switching during the evolution of small RNA pathways.

## Discussion

This study demonstrates that the function of GTSF-1 is conserved among clade V nematodes, where it associates with RRF-3 and supports its activity in the 26G-RNA pathway. In *C. elegans, C. briggsae* and *P. pacificus*, GTSF-1 uses its zinc finger region to engage a deeply conserved N-terminal domain of RRF-3 that we term the GTSF-Interacting Domain (GID), providing a mechanistic link between GTSF-1 binding and RRF-3 function. Structural predictions further support a conserved GTSF-1-RRF-3 interface at the base of nematode phylogeny, highlighting a deep evolutionary origin for this interaction. Despite extensive work on small RNA pathways, the molecular principles governing ERI complex assembly and RdRP regulation in *C. elegans* remain poorly defined, and this study begins to address that gap.

## Conserved biochemical role of GTSF-1: from PIWI* to “RRF-3*”

Across animals, GTSF1/Gtsf1 acts as a small RNA effector cofactor that converts an inactive piRNA-guided Piwi complex into a catalytically competent state (PIWI*)^16,23,24^. Biochemical reconstitution in mouse, fly, silkworm and sponge demonstrates that GTSF1 is recruited to PIWI only after proper target engagement and extensive 3′ pairing, where it induces conformational changes that activate endonucleolytic cleavage and/or factor recruitment.

In *C. elegans* and *C. briggsae*, loss of GTSF-1 collapses the RRF-3-centered ERI complex, revealing an analogous role in licensing effector function. Size-based fractionation experiments of *C. elegans* extracts indicates that GTSF-1 exists in at least two states: a smaller pre-ERIC subcomplex with RRF-3/ERI-5 and a larger mature ERIC that includes Dicer^31^, suggesting that GTSF-1 promotes stepwise assembly of the ERI complex, conceptually paralleling PIWI* maturation in other animals. By analogy with PIWI, an attractive model is that RRF-3 surveys potential RNA substrates in a low-activity state and that productive recognition, potentially defined by sequence, structure, or cofactor-marked transcripts, triggers GTSF-1 recruitment to the GID, locking RRF-3 into an active “RRF-3*” conformation competent for dsRNA synthesis and ERIC engagement. In this view, GTSF-1-dependent conformational rearrangements license dsRNA synthesis and/or productive ERIC assembly. It follows that defining what distinguishes GTSF-1-recruiting RNAs from bystander transcripts becomes a key future question, as features that recruit GTSF-1 will provide insights into substrate selection criteria for RdRP.

Similar principles may extend beyond nematodes. In the ciliate *Paramecium tetraurelia*, PtGtsf1 functions within the Polycomb Repressive Complex 2 (PRC2) to mediate Piwi-dependent transposon repression^66,67^, even though a direct Piwi-Gtsf1 interaction has not yet been demonstrated. In that context, Gtsf1 binding likely promotes PRC2 assembly or stability in a Piwi-dependent manner, reinforcing a broader view of Gtsf1 proteins as structural cofactors that stabilize or activate functional protein assemblies across deeply divergent small RNA pathways.

## PRG-1 Function Without GTSF-1: Evolutionary Loss and Functional Adaptation

All biochemically characterized PIWI proteins exhibit weak intrinsic catalytic activity, remaining in an inactive or suboptimal conformation in the absence of GTSF1^16^. In most systems, GTSF1 binding to PIWI-piRNA complexes is required either to achieve efficient target cleavage or to recruit silencing factors, suggesting that loss of GTSF1 binding would render canonical PIWI function severely compromised. This framework suggests an evolutionary scenario whereby loss of GTSF-1 binding drives repeated PRG-1 loss in nematodes: an ancestral PRG-1 that could not be productively engaged by GTSF-1 would be functionally suboptimal and, over time, dispensable.

In lineages that retain PRG-1 (clade V and early-branching Chromadorea), PRG-1 may have adopted functions that diverge from canonical PIWI activity. Studies from *C. elegans* support this hypothesis. Mutations in the DDH catalytic triad of PRG-1 do not affect piRNA levels or silencing of a piRNA sensor^25,68^, indicating that endonucleolytic cleavage is dispensable for piRNA-mediated silencing, and PRG-1 activity is coupled to RdRP (RRF-1) activity, rather than to direct target silencing. Moreover, the phenotypic consequences of PRG-1 loss are delayed rather than acutely lethal: piRNA pathway disruption leads to progressive fertility decline over multiple generations rather than immediate sterility^69^. At the molecular level*, prg-1* mutants display a striking phenotype where they accumulate aberrant 22G-RNAs on many protein-coding genes, histones, and rRNAs ^70,71^. This idea is further strengthened by the fact that 22G RNA biogenesis in absence of parentally inherited 22G RNAs can trigger acute sterility in absence of PRG-1 ^72,73^. Based on this, a currently proposed role for PRG-1 is to maintain germline homeostasis by restraining runaway small-RNA amplification. Together, these observations support a model in which clade V PRG-1 has gained GTSF-1 independent functions to act as a regulator of a small RNA network, rather than target cleavage or heterochromatin assembly.

## RdRP-GTSF1 interactions beyond nematodes

Eukaryotic “cellular” RdRPs form a distinct family from viral RdRPs and are frequently embedded in RNAi pathway ^74^, yet the molecular determinants of substrate selection and their mechanisms of action remain largely unresolved. This study reveals a core function of GTSF-1 in facilitating nematode RdRP activity, specifically for RRF-3 and raises the broader hypothesis that GTSF-1-like proteins may regulate RdRPs across eukaryotes rather than exclusively acting with PIWI proteins. Beyond nematodes, multiple metazoan lineages, including cnidarians, lophotrochozoans, and chelicerates, retain RdRP genes^10,11,74–77^. These provide rich comparative systems to test whether support for RdRP activity by GTSF-1-like proteins is conserved, or whether, in some lineages, a single GTSF-1-like protein may interface with both PIWI and RdRP complexes.

Studies from plants support the role of GTSF-1 proteins as an RdRP activator. The N-terminal region of *Zea mais* (ZmRDR2) and *Arabidopsis thaliana* (AtRDR2), that is structurally homologous to parts of GID, forms a positively charged, funnel-shaped RNA-guiding channel that captures single-stranded RNA templates and directs their 3′ ends into the catalytic cleft^62–64,78^. Cryo-EM analysis of ZmRDR2 shows that RNA/NTP binding widens the cleft angles and displaces the catalytic barrel, indicating activation through N-terminal domain motion ^64^. Mutations in these regions abolish activity without disrupting the protein’s overall fold, suggesting these domains function chiefly as conformational switches. While orthologs of GTSF-1 are only found in animals^21,79^, plants may harbor GTSF-1-like proteins that activate RdRPs by engaging their N-terminal domains through mechanisms analogous to metazoan Gtsf1-PIWI interactions.

Intriguingly, the requirement for GTSF-1 is not uniform across *C. elegans* RdRPs. RRF-1 and EGO-1 function independently of GTSF-1, generating dispersed 22G-RNAs across full-length transcripts^35,51,80,81^. By contrast, RRF-3 is thought to produce extended dsRNA precursors that are subsequently processed by Dicer^32,55,59^, a mechanism reminiscent of RdRP-Dicer interactions observed in plants and ciliates^82–85^. This suggests that long dsRNA production may impose additional requirements for processivity, substrate stabilization, or coordination with downstream processing machinery, thereby explaining why RRF-3, but not RRF-1 or EGO-1, depends on GTSF-1. Testing this model will require reconstituting RRF-3 activity with and without GTSF-1 and directly probing effects on processivity and dsRNA product length. An alternative, not mutually exclusive, possibility is that other RdRPs also rely on GTSF1-like zinc finger cofactors, but with distinct specificities. Both RRF-1 and EGO-1 harbor GID-like domains, and structural predictions strongly support their binding to GTSF-1. This raises the possibility that RRF-1 and EGO-1 are also activated by as-yet-unidentified zinc finger proteins that parallel GTSF-1 in domain architecture but diverge in sequence.

More broadly, proteins with tandem zinc finger arrays similar to GTSF-1 may partner with diverse eukaryotic RdRPs to promote complex assembly, substrate engagement, or conformational activation through conserved structural principles. Systematic surveys of physical interactions between RdRP and GTSF1-like factors, coupled with functional assays in these taxa, will be critical to distinguish between a nematode-specific innovation and a more ancient eukaryotic mechanism.

Taken together, this work shows how RNAi pathway components can diverge mechanistically while preserving essential regulatory functions. The diversification of GTSF-1 binding partners between PIWI and RdRP may be adaptive-enabling specialization of small RNA pathways in individual lineages-or neutral, arising from genetic drift without immediate advantage. This reflects evolving views that the complexity of RNAi pathways results from both selection for defense functions and constructive neutral processes^86^. In summary, this work provides important evolutionary insights into the adaptability and deep evolutionary plasticity of small RNA-mediated gene silencing systems.

## Limitations of our study

While our study provides strong support for the GTSF-1 interaction site in RRF-3 through structural predictions and genetic analyses, direct high-resolution structural validation of the GTSF-1-RRF-3 complex is still lacking. Additionally, our functional assays were performed primarily on clade V nematodes and a few related species, which limits the generalizability of our evolutionary conclusions. Lastly, although we propose mechanistic models for how GTSF-1 activates RRF-3 and licenses ERI complex assembly, these remain speculative in the absence of direct biochemical reconstitution experiments dissecting GTSF-1’s effects on RdRP enzymatic activity and ERI complex stability. Addressing these limitations will be important for solidifying our mechanistic understanding and evolutionary inferences.

## Methods

### Nematode growth and culture

Three wild-type nematode strains were used: *C. elegans* N2 (Bristol), *C. briggsae* AF16 ^87^ (Ahmedabad, India), and *P. pacificus* PS312 ^88^ (Pasadena, California). AF16 was obtained from the Caenorhabditis Genetics Center, and PS312 was provided by the Sommer Lab. All strains were maintained at 20°C on NGM plates seeded with *E. coli* OP50 under standard *C. elegans* culture conditions ^89^. Strains are listed in **Table 1**.

## Genome Editing

For *C. elegans* and *C. briggsae*, genome editing was performed by either plasmid-encoded Cas9 injections or Cas9 protein injection. For the former, protospacer sequences were identified using CRISPOR (http://crispor.tefor.net) ^90^ and inserted into pRK2411 (a derivative of pDD162, Addgene #47549) through site-directed, ligase-independent mutagenesis (SLIM) ^91^. SLIM products were introduced into DH5α competent cells (Invitrogen) and grown on LB agar containing 100 μg/ml ampicillin and purified. Injection mixes contained 50 ng/µl Cas9 + co-conversion sgRNA plasmid, 50 ng/µl “gene of interest” sgRNA plasmid, 750 nM co-conversion ssODN donor Integrated DNA Technologies™ (IDT) and 750 nM gene of interest ssODN donor (IDT). For *Cel-gtsf-1(xf264)*, 400 ng/µl PCR product was used as a linear, double-stranded DNA donor template. For Cas9 protein injections, Alt-R CRISPR-Cas9 tracrRNA and Alt-R CRISPR-Cas9 Custom Guide RNAs were obtained from IDT and the Predesigned Alt-R™ CRISPR-Cas9 guide RNA tool was used to determine protospacer sequences. Strains were created by microinjecting recombinant Cas9 protein (prepared in-house), homologous recombination templates (IDT), and guide RNA molecules (IDT) as described previously ^92^. Phenotypes generated through *dpy-10(cn64)* ^93^ and crRNA against *dpy-1* in *C. elegans* and *C. briggsae* respectively, served as co-conversion markers. The *P. pacificus gtsf-1* mutant was generated using established CRISPR-Cas9 protocols with target-specific crRNA, universal tracrRNA (IDT ALT-R), and Cas9 protein (IDT Cat.#1081058;^94,95^). crRNAs were designed to target sequences 20 bp upstream of protospacer adjacent motifs and verified against the El Paco genome assembly ^96^ to exclude off-target sites. For all species, mixes were microinjected into both gonad arms of 10–20 young adult hermaphrodites at 20°C. F1 progeny were screened by PCR, and edits were confirmed by Sanger sequencing. All strains were outcrossed at least twice before analysis. Guide sequences and repair templates are provided in **Table 2**.

## Brood size calculation

Animals synchronized by hypochlorite treatment were maintained on OP50-seeded NGM plates at 20°C until the early L3 larval stage. Twenty to thirty individual L3 worms per strain were then singled and incubated at either 20°C or 25-30°C. Adult hermaphrodites were transferred daily to fresh plates for five consecutive days or until egg-laying ceased. Progeny were counted 48 hours post-transfer for assays at 20°C (to facilitate accurate counting at the L3-L4 stage) or 24 hours post-transfer for assays at 25-30°C. Total brood size was calculated as the sum of all viable progeny per hermaphrodite. Data are presented as mean brood size per strain. Statistical comparisons between mutant and wild-type strains were performed using the Wilcoxon rank-sum test with Benjamini-Hochberg correction in RStudio.

## RNA interference

RNAi experiments were conducted as previously described^97,98^. For *C. elegans dpy-13* RNAi, synchronized L1 larvae were seeded on six-well plates and scored for dumpy phenotype on the second day of adulthood.

## CbrGTSF-1 and PpaGTSF-1 antibody production

### a. Recombinant GTSF-1 orthologs protein production

pET vectors containing coding sequences for tagged, full length GTSF-1 orthologs from *C. briggsae* and *P. pacificus* (His_6_-MBP-TEVsite-GTSF-1 and His_6_-SMT3-3Csite-GTSF-1) were transformed into *E. coli* (BL21-CodonPlus (DE3)-RIL, Agilent). Cells were grown in 2 liters of LB Broth Miller (Formedium), containing 50µg/ml Kanamycin at 37°C/ 140 rpm until OD(600) of 0.6. Expression was induced by addition of 0.5 mM IPTG (neoFroxx) and further incubation at 18°C overnight. Cells were harvested by centrifugation (4k x g, 15 min, 4°C), re-suspended in ice cold lysis buffer (50 mM Tris-Cl, 500 mM NaCl, 15 mM imidazole, 1 mM MgCl2, 0.5 mM DTT, 1x Roche cOmplete EDTA-free protease inhibitor, 50 U/ml Sigma Benzonase, 0.1% Triton X-100, pH 8.0) and lysed by sonication at 4°C (Branson Sonifier 450, 9 mm tip, output 7, 20% duty cycle, 2 x 3 min). Cell lysates were cleared by centrifugation (40k x g, 30 min, 4°C) and applied to a HisTrap FF 5 ml column (Cytiva) at 4°C, using an automated chromatography system (Biorad NGC Quest Plus). The column was washed with 20 CV (column volumes) of wash buffer (50 mM Tris-Cl, 500 mM NaCl, 15 mM imidazole, 5 % glycerol). Proteins were eluted from the column by applying a linear gradient of 15-500 mM imidazole (in 50 mM Tris-Cl, 500 mM NaCl, 5% glycerol) over 10 CV and peak elution fractions were pooled for further purification. For the purification of untagged GTSF-1 orthologs, 25 to 50 mg of each His_6_-MBP-TEV-GTSF-1 ortholog was digested using 1 mg recombinant His_6_-TEV protease at 4°C overnight during a 1:100 (v/v) dialysis (50 mM Tris-Cl, 200 mM NaCl, 0.5 mM DTT, 5 % glycerol), using SnakeSkin dialysis tubing with 3.5 kDa cut-off (Thermo Fisher Scientific).

The dialysed solution, containing untagged GSTF-1, His6-MBP-TEV and His_6_-TEV protease, was supplemented with 15 mM imidazole (pH 8.0) and applied to a HisTrap FF 5 ml column. The flow through containing untagged GTSF-1 was collected for further purification. Untagged and His_6_-SMT3-3C-tagged GTSF-1 orthologs were concentrated using Amicon Ultra-15 spin concentrators with 3 kDa cut-off (Merck Millipore) and subjected to size exclusion chromatography at 4°C (Superdex 75 16/60 pg, Cytiva, in sterile PBS). Peak elution fractions of the proteins were analysed by SDS-PAGE, pooled, spin-concentrated to a range of 1.5 to 4 g/l, aliquoted, flash-frozen in liquid nitrogen and stored at -80°C. Untagged and SMT-3-tagged *P. pacificus* GTSF-1 was oligomeric and eluted in the void volume of the Superdex 75 column.

### **b.** Polyclonal anti-GTSF-1 orthologs antibody production

Polyclonal antibodies against recombinant, untagged *C. briggsae* and *P. pacificus* GTSF-1 were raised in rabbit, using Eurogentećs Speedy 28-day program. 200 µg of GTSF-1 in PBS were used for each immunization round. For antibody purification from serum, 2 to 3 mg of each His_6_-SMT3-3Csite-tagged GTSF-1 ortholog was diluted with 4 ml coupling buffer (50 mM Tris, 5 mM EDTA, pH 8.5) and incubated with 2 ml SulfoLink bead slurry (Thermo Fisher Scientific), pre-equilibrated in coupling buffer. The proteins were covalently coupled to beads by incubation at room temperature for 2 h in a Poly-Prep column (Bio-Rad) while rotating. After washing with coupling buffer, the remaining iodoacetyl groups were blocked with 5 ml quenching buffer (50 mM L-cysteine in coupling buffer) for 30 min. The resin was subsequently washed with 10 ml coupling buffer containing 1 M NaCl, followed by 20 ml PBS. For antibody purification, 5 ml rabbit serum was diluted 1:2 with PBS, filtered through a 0.45 μm filter and incubated with the respective GTSF-1-coupled SulfoLink resin in a Poly-Prep column overnight at 4°C while rotating. Resins were washed with 10 ml PBS containing 0.1% Triton X-100 followed by 20 ml PBS. Antibodies were eluted with 6 x 1 ml low pH buffer (200 mM glycine-HCl, 150 mM NaCl, pH 2.3) and immediately neutralized with 0.2 ml of 1 M Tris-Cl pH 8.5. Eluted fractions were analysed by SDS-PAGE and rebuffered into storage buffer (PBS, 10% glycerol, 0.05% NaN_3_) using PD-10 columns (Cytiva). Antibodies were spin-concentrated to 1 g/l, aliquoted, flash-frozen in liquid nitrogen and stored at -80°C.

## Immunoprecipitations

### a. IP with anti-CbrGTSF-1, anti-PpaGTSF-1 and anti-IgG antibodies

Animals were expanded on OP50 plates for two generations, following which L1 larvae were synchronized by hypochlorite treatment and seeded onto high-density OP50 egg plates. 76 hours post-seeding, gravid adults were harvested by snap-freezing in liquid nitrogen (LN₂) and stored at -80°C. For IPs, 200 µl of gravid adult samples were lysed in an equal volume of 2x lysis buffer (50 mM Tris-HCl pH 7.5, 300 mM NaCl, 3 mM MgCl₂, 2 mM DTT, 0.2% Triton X-100, and cOmplete Mini EDTA-free Protease Inhibitor Cocktail). Lysates were sonicated using a Bioruptor Plus at 4°C for 10 cycles (30 seconds ON/30 seconds OFF), then centrifuged at 21,000 ×g for 10 minutes at 4°C. Protein concentrations in the soluble lysate were measured using the Pierce BCA Protein Assay and normalized to 1–3 mg total protein with 1x lysis buffer. Lysates were pre-cleared by incubation with 30 µl DYNAL Dynabeads™ Protein G (equilibrated in lysis buffer) for 1–2 hours at 4°C. For each immunoprecipitation, 30 µl Dynabeads™ Protein G were washed three times with 1x wash buffer (25 mM Tris-HCl pH 7.5, 150 mM NaCl, 1.5 mM MgCl₂, 1 mM DTT, and cOmplete Mini EDTA-free Protease Inhibitor Cocktail), then conjugated to 1.5–2 µg anti-CbrGTSF-1, anti-PpaGTSF-1 (in-house) or anti-IgG antibody according to manufacturer’s instructions. Pre-cleared lysates were incubated with antibody-conjugated Dynabeads™ for 2 hours at 4°C with rotation. Beads were washed three times with wash buffer, and bound proteins were eluted in NuPAGE LDS sample buffer supplemented with 100 mM DTT by heating at 70°C for 10 minutes. Eluted samples were analyzed by mass spectrometry or western blot.

### **b.** IP with anti-FLAG and anti-MYC antibodies

For embryo collection, synchronized L1 larvae were seeded onto 15 cm OP50 plates and harvested at 96 hours by hypochlorite treatment. Embryos were washed extensively with cold M9, resuspended in 1x lysis buffer (25 mM Tris-HCl pH 7.0, 150 mM NaCl, 1 mM DTT, 1.5 mM MgCl₂, 0.1% Triton X-100, 1× cOmplete Mini Protease Inhibitor), and snap-frozen as droplets in liquid nitrogen for storage at −80°C. Frozen droplets were ground to a fine powder in a liquid nitrogen-cooled mortar, then further homogenized with 40 strokes using a chilled Dounce homogenizer (pestle B). Lysates were clarified by centrifugation at 21,000×g for 10 minutes at 4°C, and protein concentrations in the supernatant were normalized by BCA assay. For immunoprecipitation, 30 µg *C. briggsae* lysate was incubated with 10 µL anti-cMYC magnetic beads (Sigma) to capture 3xMYC::RRF-3, or 100 µg *C. elegans* lysate was incubated with 15 µL anti-FLAG M2 magnetic beads (Sigma) to capture 3xFLAG::RRF-3; both reactions incubated overnight at 4°C with rotation. Beads were magnetically separated and washed three times with wash buffer (25 mM Tris-HCl pH 7.0, 150 mM NaCl, 1 mM DTT, 1.5 mM MgCl₂, 1× cOmplete Mini Protease Inhibitor). Proteins were eluted by heating beads in 25 mM Tris-HCl pH 8.0 containing 1% SDS at 70°C for 10 minutes, and eluates were stored at −20°C for downstream processing.

## Mass-spectrometry

### a. Enzymatic protein digestion

For mass spectrometry analysis, immunoprecipitated samples were processed using two different protocols. For GTSF-1 immunoprecipitation in *C. briggsae* and *P. pacificus* (SetA), the samples were separated on a 4–12% NOVEX NuPAGE gradient SDS gel (Thermo Fisher Scientific) for 10 min at 180 V in 1× MES buffer (Thermo Fisher Scientific). Proteins were fixed and stained with Coomassie G250 Brilliant Blue (Carl Roth). The gel lanes were cut, minced into pieces and transferred to an Eppendorf tube. Gel pieces were destained with a 50% ethanol/50 mM ammonium bicarbonate (ABC) solution. Proteins were reduced in 10 mM DTT (Sigma-Aldrich) for 1 h at 56 °C and then alkylated with 5 mM iodoacetamide (Sigma-Aldrich) for 45 min at room temperature. Proteins were digested with trypsin (Sigma-Aldrich) overnight at 37 °C. Peptides were extracted from the gel by two incubations with 30% ABC/acetonitrile and three subsequent incubations with pure acetonitrile. The acetonitrile was subsequently evaporated in a concentrator (Eppendorf) and loaded onto StageTips^99^ for desalting and storage.

For RRF-3 immunoprecipitation in *C. briggsae* and RRF-3/GTSF-1 dual immunoprecipitation in *C. elegans* (Set B), the eluted proteins were processed using the SP3 approach^100^. The proteins were then digested using trypsin overnight at 37°C. The resultant peptide solution was purified in C_18_ StageTips ^101^.

For Set B samples, peptides were separated in an in-house packed 30-cm analytical column (inner diameter: 75 μm; ReproSil-Pur 120 C_18_-AQ 1.9-μm beads, Dr. Maisch GmbH) by online reverse phase chromatography on an EASY-nLC 1000 UHPLC system (Thermo Scientific) through a 105-min non-linear gradient of 1.6-32% acetonitrile with 0.1% formic acid at a nanoflow rate of 225 nl/min. The eluted peptides were sprayed directly by electrospray ionization into a Q Exactive Plus Orbitrap mass spectrometer (Thermo Scientific).

### **b.** Liquid chromatography tandem mass spectrometry

Peptides from both sample sets were separated by online reverse-phase chromatography on an EASY-nLC 1000 UHPLC system (Thermo Scientific). For Set A samples, peptides were separated on a 20 cm in-house packed analytical column and for Set B a 30cm in-house packed column (inner diameter: 75 μm; ReproSil-Pur 120 C_18_-AQ 1.9-μm beads, Dr. Maisch GmbH). For set A We used a 94 min gradient from 2 to 40% acetonitrile and for Set B a 105-min non-linear gradient of 1.6-32% acetonitrile, both in 0.1% formic acid at a flow rate of 225 nl min^−1^. The eluted peptides were sprayed directly by electrospray ionization into a Q Exactive Plus Orbitrap mass spectrometer (Thermo Scientific).

For Set A, the mass spectrometer was operated with a top 10 MS/MS data-dependent acquisition scheme per MS full scan. For Set B, mass spectrometry measurement was conducted in data-dependent acquisition mode using a top10 method with one full scan (mass range: 300 to 1,650 m/z; resolution: 70,000, target value: 3 × 10^6^, maximum injection time: 20 ms) followed by 10 fragmentation scans via higher energy collision dissociation (HCD; normalised collision energy: 25%, resolution: 17,500, target value: 1 × 10^5^, maximum injection time: 120 ms, isolation window: 1.8 m/z). Precursor ions of unassigned or +1 charge state were rejected. Additionally, precursor ions already isolated for fragmentation were dynamically excluded for 20 s.

### **c.** Mass spectrometry data processing and statistical analysis

For all samples, mass spectrometry raw data were processed by MaxQuant software package (version 2.1.3.0)^102^ using its built-in Andromeda search engine^103^. Spectral data were searched against a target-decoy database consisting of the forward and reverse sequences of the UniProt *C. elegans* (August 2014; 27,814 entries), UniProt *C. briggsae* (release 2023_02; 21,756 entries) and UniProt *P. pacificus* (release 2023_03; 26,122 entries) reference proteome, the UniProt *E. coli* (release 2023_01; 5,064 entries) reference proteome and a list of common contaminants. Trypsin/P specificity was assigned. Carbamidomethylation of cysteine was set as fixed modification. Methionine oxidation and protein N-terminal acetylation were chosen as variable modifications. A maximum of 2 missed cleavages were tolerated. The “second peptides” options were switched on. The “match between runs” function was activated. The minimum peptide length was set to be 7 amino acids. False discovery rate (FDR) was set to 1% at both peptide and protein levels.

The MaxLFQ algorithm^104^ was employed for label-free protein quantification without using its default normalization option. Minimum LFQ ratio count was set to be one. Both the unique and razor peptides were used for quantification. Detected *E. coli* proteins, reverse hits, potential contaminants and “only identified by site” protein groups were filtered out. Data normalization was performed on the log-transformed data using a median-centering approach. Proteins were further filtered to retain only those detected in at least two out of the four replicates in either group of each comparison.

### **d.** Statistical analysis

For Set A, the label-free quantification values were log_2_-transformed and the median across the replicates was calculated. This enrichment was plotted against the −log_10_-transformed *P* value (Welch’s *t*-test) using the ggplot2 package in the R environment.

For Set B, following imputation of the missing LFQ intensity values, the statistical significance of the difference between the two groups was assessed using a modified *t*-statistic (*t*(SAM, statistical analysis of microarrays)^105^ and visualized on a volcano plot.

## GST pulldown assay

### a. GST-on bead purification

Sequences of *C. elegans* gtsf-1 (wild-type or mutant) and *B. mori* gtsf-1 were amplified from cDNA and cloned into pGEX-6P-1 (Addgene) using Gibson assembly ^106,107^. Plasmids were transformed into *E. coli* BL21 DE3 Rosetta cells and grown in 75 mL LB Broth containing 100 μg/mL ampicillin at 37°C/140 rpm to OD₆₀₀ = 0.6. Expression was induced with 0.5 mM IPTG (Promega) and cultures incubated overnight at 18°C. Cells were harvested by centrifugation (4,000×g, 20 min, 4°C) and resuspended in 5 mL ice-cold lysis buffer (50 mM Tris-HCl pH 7.5, 150 mM NaCl, 1 mM DTT, 0.1% Triton X-100, 1 μg/μL lysozyme, 1×cOmplete EDTA-free protease inhibitor). Lysates were sonicated at 4°C (Branson Sonifier 450, 5 mm tip, output 5, 15% duty cycle, 2×4 min), then cleared by centrifugation (4,000×g, 30 min, 4°C). Cleared lysates were incubated with 10 μL Glutathione High-Capacity Magnetic Agarose Beads (Sigma) overnight at 18°C with end-to-end mixing. GST-GTSF-1 beads were washed 3 times with 500 μL wash buffer (50 mM Tris-HCl pH 7.5, 150 mM NaCl, 1 mM DTT, 1 mM MgCl₂, 1×cOmplete EDTA-free protease inhibitor). For loading control, 10% of beads were eluted by heating at 70°C in 1×NuPAGE™ LDS sample buffer (Invitrogen) containing 100 mM DTT. Samples were centrifuged (21,000×g, 5 min) and supernatants loaded onto NuPAGE™ Bis-Tris 4–12% gradient gels and run in MOPS buffer at 200 V for 50 min. GST-GTSF-1 proteins were visualized by InstantBlue™ staining.

### **b.** GST pulldown

Remaining GST-GTSF-1 beads were incubated with *C. elegans* RFK421(*rrf-3* (*pk1426*) II; *xfIs63*[*gld-1*(*prm*)::*flag*::*rrf-3*::*unc-54* (*3’UTR*)] IV; *gtsf-1*(*xf43*) IV) embryo lysate in 1x lysis buffer (25 mM Tris-HCl pH 7.0, 150 mM NaCl, 1 mM DTT, 1.5 mM MgCl₂, 0.1% Triton X-100, 1× cOmplete Mini Protease Inhibitor) for 2 h at 4°C with end-to-end mixing. As a positive control, 5 μL Anti-FLAG® M2 Magnetic beads (Sigma) were washed 3 times with embryo lysis buffer and incubated with 15% of the embryo lysate alongside GST-GTSF-1 beads. All beads were washed 3 times with 500 μL wash buffer (25 mM Tris-HCl pH 7.5, 150 mM NaCl, 1 mM DTT, 1.5 mM MgCl₂, 1×cOmplete Mini Protease Inhibitor). Proteins were eluted by heating at 70°C in 1×NuPAGE™ LDS sample buffer containing 100 mM DTT, centrifuged (21,000×g, 5 min), and supernatants loaded onto NuPAGE™ 3–8% Tris-Acetate gels and run in Tris-Acetate buffer at 150 V for 1 h. 3×FLAG::RRF-3 was visualized by Western blot as described below.

## Western Blot

Western blotting was performed on lysates from whole animal lysates, sonicated or liquid nitrogen homogenized samples. For whole animal, 100 gravid adult hermaphrodites were hand-picked and collected directly into 1× NuPAGE™ LDS sample buffer (Invitrogen) containing 100 mM DTT and heated at 95°C for 30 min before clarification by centrifugation (21,000×g, 10 min) and storage at −20°C. Sonicated or liquid nitrogen homogenized samples were prepared as described in previous sections. For detection of 3×FLAG::RRF-3, samples were separated on NuPAGE™ 3–8% Tris-Acetate gels in Tris-Acetate running buffer at 150 V for 1 h and for GTSF-1 and GTSF-1::3×FLAG detection, samples were separated on NuPAGE™ 10% or 12% Bis-Tris gels in MES running buffer at 200 V for 35 min, alongside Color Prestained Protein Standard (10–250 kDa, NEB P7719S). Proteins were transferred to 0.45 µm nitrocellulose membrane (Amersham Protan GE10600002) in NuPAGE™ Transfer Buffer with 10% methanol at 20 V overnight at 4° C, and membranes were blocked in 5% skim milk in PBS-T (0.1% Tween-20) for 30 min. Blocked membranes were sectioned by molecular weight and incubated for 1h at room temperature with primary antibodies in blocking buffer (1:500 rabbit anti-CelGTSF-1 (Q5963 ^31^); 1:1000 rabbit anti-FLAG (Sigma-Aldrich F7425-2MG); 1:1000 mouse anti-FLAG (Sigma-Aldrich F3165); 1:1000 mouse anti-MYC (CST 2276S); 1:1000 mouse anti-α-tubulin (Abcam ab7291), followed by three 10-min washes in PBS-T. Membranes were then incubated for 1 h with HRP-conjugated anti-rabbit or anti-mouse secondary antibodies (CST 7074S and 7076S), washed three times, and developed using SuperSignal™ West Atto Femto Maximum Sensitivity substrate or SuperSignal™ West Atto Ultimate Sensitivity substrate (for GTSF-1) or (Thermo Scientific) according to the manufacturer’s instructions.

## Yeast Two-Hybrid

Genes of interest were amplified from cDNA, and point mutations were introduced using the SLIM protocol^91^. Constructs were cloned into pGAD and pGBD plasmids as described previously^108^. Double plasmid transformation was performed in *Saccharomyces cerevisiae* strain AH109^108^ using the high-efficiency LiAc method^109^. Briefly, cells were grown to mid-log phase (OD₆₀₀ 0.55–0.7), washed, and resuspended in 1× TE/LiOAc before being mixed with 0.15 µg each of AD and BD plasmids and 0.1 µg denatured UltraPure™ Salmon Sperm DNA (Invitrogen, 15632011). Following incubation with PEG/TE/LiOAc solution (30 °C, 30 min), cells were heat-shocked at 42 °C for 15 min with DMSO, plated on selective media supplemented with adenine and histidine, and incubated for 3 days at 30 °C. Transformed colonies were resuspended in sterile water, and OD₆₀₀ was measured using an Infinite® M200 Pro plate reader. Cells were diluted to 5×10⁶ and 5×10⁵ cells/mL (assuming OD₆₀₀ 0.7 = 1×10⁷ cells/mL), and 5 µL of each dilution was spotted onto control plates or selection plates lacking adenine only or both adenine and histidine. Plates were imaged after 3 days of incubation at 30 °C.

All selective media were prepared using Yeast Synthetic Drop-out Medium Supplements without histidine, leucine, tryptophan, and adenine (Sigma-Aldrich, Y2021) and Yeast Nitrogen Base Without Amino Acids (Sigma-Aldrich, Y0626) at final concentrations of 1.399 g/L and 6.7 g/L, respectively. Adenine (Sigma-Aldrich, A2786) and L-histidine monohydrochloride monohydrate (Sigma-Aldrich, H5659) were added to supplemented plates at 21 mg/L and 85.6 mg/L, respectively. Bacto Agar (Sigma-Aldrich, A5306) was included at 20 g/L, and all media were supplemented with 2% glucose.

## RNA extraction, sequencing and analysis

### a. RNA extraction

Synchronized animals were cultured at 20°C and harvested at gravid-adult or late-L4 stages. Embryo samples were collected from hypochlorite treated synchronized gravid adults. All samples were collected in quadruplicates, snap-frozen in 50 µL pellets on dry ice and stored at - 80°C until RNA extraction. For RNA extraction, pellets were resuspended in 500 µL TRIzol™ LS Reagent (Invitrogen, 10296028) and subjected to six freeze-thaw cycles (liquid nitrogen for 30 s, 37°C water bath for 1 min). Samples were vortexed between cycles to ensure complete lysis and verified microscopically by the absence of intact worms. Debris were clarified by centrifugation at 21,000×g for 5 min. The supernatant was mixed with 100% ethanol (1:1 v/v) and processed immediately Direct-zol™ RNA Miniprep kit (Zymo Research) according to the manufacturer’s instructions.

### **b.** Library preparation and sequencing

RppH Treatment: 200 to 1000 ng RNA was subjected to RNA 5’ Pyrophosphohydrolase (RppH) treatment (NEB, M0356S) at 37°C for 1 hour to eliminate pyrophosphate from the 5’ end of triphosphorylated RNA, leaving a 5’ monophosphate RNA. Following purification, the samples were quantified using the Qubit RNA High Sensitivity Assay Kit.

NGS Library Preparation and Sequencing: Next-generation sequencing (NGS) library preparation employed the NEXTflex Small RNA-Seq Kit V3, following Bioo Scientific’s standard protocol (V19.01) from Step A to Step G. The process utilized the NEXTFlex 3’ SR Adaptor and 5’ SR Adaptor (5’rApp/NNNNTGGAATTCTCGGGTGCCAAGG/3ddC/ and 5’GUUCAGAGUUCUACAGUCCGACGAUCNNNN, respectively). Step A (NEXTflex 3’4N Adenylated Adapter Ligation) was conducted overnight at 20°C.

Libraries were prepared with an initial amount of 500 to 900 ng and amplified through 14 to 15 PCR cycles. Amplified libraries underwent purification via an 8% TBE gel, and fragments in the 15–40 nt range were size-selected. Profiling occurred using a High Sensitivity DNA Chip on a 2100 Bioanalyzer (Agilent Technologies), and quantification was performed with the Qubit dsDNA HS Assay Kit on a Qubit 2.0 Fluorometer (Life Technologies). Samples were equimolarly pooled and sequenced on a Highoutput NextSeq 500/550 Flowcell, single-read for 1x 84 cycles, plus 7 cycles for the index read.

## **c.** Read processing and mapping

Analysis of small-RNA sequencing datasets was performed using TinyRNA^110^. Briefly, the preprocessing of FASTQ files, with adapter trimming and quality filtering, was executed through the utilization of fastp^111^. (-a TGGAATTCTCGGGTGCCAAGG, -q 20 -e 25 –u 10). 5’ and 3’ random UMIs (NNNN-RNA sequence-NNNN) were then removed and reads shorter than 15 nucleotides were filtered out. Subsequently, unique sequences were counted and collapsed using tiny-collapse. Collapsed reads were aligned to *C. elegans* (PRJNA13758, Worm base version WS279), *C. briggsae* (PRJNA10731, WS285) and *P. pacificus* (PRJNA12644, El Paco assembly) reference genome using bowtie (v.1.3.1)^112^. Features were assigned using tiny-count^110,113^. Feature assignment is based on features annotated in custom GFF input files. Differential expression analysis is performed by tiny-deseq, which uses DESeq2^114^.

Feature annotation: Custom GFF3 annotation files were generated for each nematode species to enable accurate feature-based quantification of small RNA reads. For *Caenorhabditis elegans*, annotations from WormBase Release WS279 (c_elegans.PRJNA13758.WS279.annotations.gff3.gz) containing major transcript features (asRNA, coding_genes, lincRNA, miRNA, ncRNA, piRNA, pseudogene, rRNA, scRNA, snoRNA, snRNA, transposons, tRNA) were merged with mature miRNA loci from miRBase v22. For *Caenorhabditis* briggsae, the WormBase WS285 annotation (c_briggsae.PRJNA10731.WS285.annotations.gff3.gz) was combined with miRBase v22 and piRNA loci from^115^. For *Pristionchus pacificus*, the coding genes from WormBase Parasite WBPS19 annotation (pristionchus_pacificus.PRJNA12644.WBPS19.annotations.gff3) was merged with miRNA coordinates from miRBase v22 and piRNA features from^115^. For both *C. briggsae* and *P. pacificus*, piRNAs that were not assembled into chromosomes were excluded from the annotation file.

Feature count rules: Small RNA reads were hierarchically assigned to biotypes using the custom merged GFF3 annotations for each species, with classification rules based on genomic overlap, strand, 5′ end nucleotide, and read length. For *C. elegans* and *C. briggsae*, reads mapping partially to annotated rRNA loci were first classified as rRNAs and excluded from downstream analyses; remaining reads mapping 5′-anchored in sense orientation to annotated miRNA loci were classified as miRNAs, followed by sense 5′-anchored reads of length 18–21 nt at piRNA loci as piRNAs, antisense partially overlapping 26-nt 5′G gene-derived reads as 26G-RNAs, and antisense partially overlapping 21–23-nt 5′G gene-derived reads as 22G-RNAs. In contrast, for *P. pacificus* the same hierarchical scheme for miRNAs, piRNAs, 26G-RNAs, and 22G-RNAs was applied directly without prior removal of rRNA-mapping reads.

Counts were normalized with DESeq2 across three biological replicates for each genotype. For all small RNA classes, genes were required to have a mean of at least 5 normalized counts across the three wild-type replicates to be retained for analysis. Relative abundance was calculated as the mean normalized counts in mutant samples divided by the sum of the mean normalized counts in mutant and wild-type samples. Genes were classified as 26G-RNA targets if they were assigned to the 26G-RNA class and showed a DESeq2-adjusted P value (padj) < 0.05 and a log2 fold change < −2 in mutants compared with wild type.

## AlphaFold predictions

*C. elegans*, *C. briggsae* and *P. pacificus* protein sequences were retrieved from WormBase. Monomer and multimer structure predictions were performed using AlphaFold3 ^116^ (https://alphafoldserver.com), with 2 copies of Zn^2+^ where specified. To predict potential protein–protein interactions via high-throughput screening, amino acid sequences were retrieved from UniProt or reciprocal BLAST and formatted as FASTA files. Two sequence sets were generated: a bait and a prey file **(Table 2).** All pairwise combinations between bait and prey sequences were systematically screened using the ht-colabfold pipeline (https://gitlab.com/BrenneckeLab/ht-colabfold) as previously described^117^, which automates large-scale structure prediction with AlphaFold-Multimer (v2)^118–120^ via ColabFold^5^. Predictions were conducted at the Life Science Compute Cluster (LiSC) of the University of Vienna, using default parameters, including multiple sequence alignments generated via MMseqs^118^. Structural models were ranked based on ipTM and PEAK scores ^117^.

## ConSurf Analysis

Conservation analysis of GTSF-1 was performed using orthologous sequences from 26 nematode species: *C. elegans*, *C. briggsae*, *C. nigoni*, *C. brenneri*, *C. tropicalis*, *C. doughertyi*, *C. inopinata*, *C. kamaaina*, *C. japonica*, *C. plicata*, *C. virilis*, *C. castelli*, *C. nouraguensis*, *C. sp21*, *C. sp26*, *C. sp28*, *C. sp29*, *C. sp31*, *C. sp38*, *C. sp39*, *C. sp40*, *P. pacificus*, *Toxocara canis*, *Wuchereria bancrofti*, *Brugia malayi*, and *Brugia pahangi*. Orthologous RRF-3 protein sequences were retrieved from WormBase for 11 nematode species: *C. elegans*, *C. briggsae*, *C. nigoni*, *C. remanei*, *C. latens*, *C. brenneri*, *C. angaria*, *C. bovis*, *C. japonica*, *C. sp36*, and *P. pacificus*. Sequences were obtained from UniProt through BLAST using *C. elegans* GTSF-1 as reference. Multiple sequence alignment was generated using Clustal Omega (https://www.ebi.ac.uk/jdispatcher/msa/clustalo) and submitted to ConSurf server (https://consurf.tau.ac.il/)^121^ along with the top-ranking AlphaFold3 prediction of GTSF-1–2×Zn²⁺ monomer for GTSF-1 scoring or GTSF-1–RRF-3–2×Zn²⁺ complex for RRF-3 scoring. ConSurf assigned conservation grades on a 1–9 scale (1 = variable, 9 = highly conserved) to each residue. The output structure file with mapped conservation scores was visualized and analyzed in ChimeraX.

## Genome data

Protein coding gene sets from 69 publicly available nematode genomes were analyzed, representing all five major nematode clades (I-V). To provide an outgroup, four genomes from the sister phylum Nematomorpha - *Acutogordius australiensis*, *Nectonema munidae*, *Gordionus montsenyensis* and *Gordius aquaticus* were additionally included **(Table 3).**

## Identification of piRNA Pathway Orthologues

To identify orthologues of the core *C. elegans* piRNA pathway proteins, we assembled a curated reference set consisting of PRG-1, GTSF-1, EGO-1, RRF-1, and RRF-3 from the *C. elegans* genome on Wormbase (WS276). Orthologues in each target species were identified using the reciprocal best BLASTP hit (RBH) approach, a widely established method for orthology inference. A candidate gene was retained as a putative one-to-one orthologue only if each sequence was the top BLASTP hit of the other under identical search parameters. This strict reciprocal criterion provided a conservative baseline for orthologue detection ^122,123^.

## Figures

Molecular graphics and analyses performed with UCSF ChimeraX^124^. Electrostatic surface calculations were performed with APBS^125^ with a solvent ion concentration of 0.15 M using the AMBER force field. The ggplot2 package of RStudio was used to produce graphs ^126^.

## Tables

Table 1: Nematode strains used in this study

Table 2: Protein sequences used for HT-AF2 screen Table 3: Summary of the species datasets

## Acknowledgements

We gratefully acknowledge support from the IMB Genomics Core Facility for RNA sequencing. We thank the IMB Media Laboratory for providing consumables and the Proteomics Core Facility for conducting mass-spectrometry experiments. We are grateful to Martin Möckle and the Protein Production Core Facility at IMB for purifying recombinant GTSF-1 orthologs and generating anti-GTSF1 polyclonal rabbit antibodies. AlphaFold high-throughput computational analyses were performed using the Life Science Compute Cluster (LiSC) at the University of Vienna. We thank Dr. Jonathan Ipsaro for valuable discussions regarding the RNA-binding properties of nematode GTSF-1. The work was funded by grants from the Deutsch Forschungsgemeinschaft: GenEvo - project number 407023052 (RFK) and the Wellcome Trust (How to make a Parasite).

## Statement on use of AI

Parts of this manuscript were edited with the AI software Claude and Perplexity.

## Data Repositories

All sequencing data generated for this study has been uploaded to the European Nucleotide Archive (www.ebi.ac.uk) under accession number PRJEB105295. All mass spectrometry data generated in this study have been uploaded to the UCSD Mass Spectrometry Interactive Virtual Environment (MassIVE) and to the PRoteomics IDEntifications Database (PRIDE) server. Mass spectrometry data for GTSF-1 immunoprecipitation in *C. briggsae* and *P. pacificus* are available under PRIDE accession number PXD072041. Data for RRF-3 immunoprecipitation in *C. briggsae* and RRF-3/GTSF-1 dual immunoprecipitation in *C. elegans* are under MassIVE accession number.

## Reviewer access details

PRIDE Project accession: PXD072041 Token: m06152ky8kJG

Username: Password: N56zwlIK0V2Q

GTSF-1 in *P. pacificus*

MassIVE Accession ID: MSV000100223

#for reviewers ftp://MSV000100223@massive-ftp.ucsd.edu Username: MSV000100223_reviewer

Password: h#BpL6$MwKKE9%X

GTSF-1 and RRF-3 in *C. elegans*

MassIVE Accession ID: MSV000100222

#for reviewers ftp://MSV000100222@massive-ftp.ucsd.edu

Username: MSV000100222_reviewer Password: h#BpL6$MwKKE9%X

RRF-3 in *C. briggsae*

MassIVE Accession ID: MSV000100221

#for reviewers ftp://MSV000100221@massive-ftp.ucsd.edu Username: MSV000100221_reviewer Password: h#BpL6$MwKKE9%X

## Author contributions (CRediT taxonomy)

**Shamitha Govind:** Investigation (lead), Conceptualization (lead), Writing – original draft, Visualization (lead), Methodology (lead), Formal analysis. **Sebastian Ruppert:** Investigation (lead), Writing – review & editing (supporting), Visualization (supporting). **Joseph Kirangwa:** Investigation (supporting). **Virginia Busetto:** Formal analysis, Writing – review & editing (supporting). **Emily Nischwitz:** Formal analysis. **Miguel Almeida:** Investigation (supporting), Conceptualization (supporting), Writing – review & editing (supporting). **Svenja Hellmann:** Investigation (supporting). **Hahn Witte:** Resources. **Ralf J. Sommer:** Resources. **Falk Butter:** Formal analysis. **Sebastian Falk:** Resources, Formal analysis. **Peter Sarkies:** Investigation (supporting), Methodology (supporting), Writing – review & editing (supporting), Supervision (supporting). **René F. Ketting:** Writing – review & editing (lead), Methodology (supporting), Conceptualization (lead), Project administration, Funding acquisition, Supervision (lead).

---

**Supplementary Figure 1.**
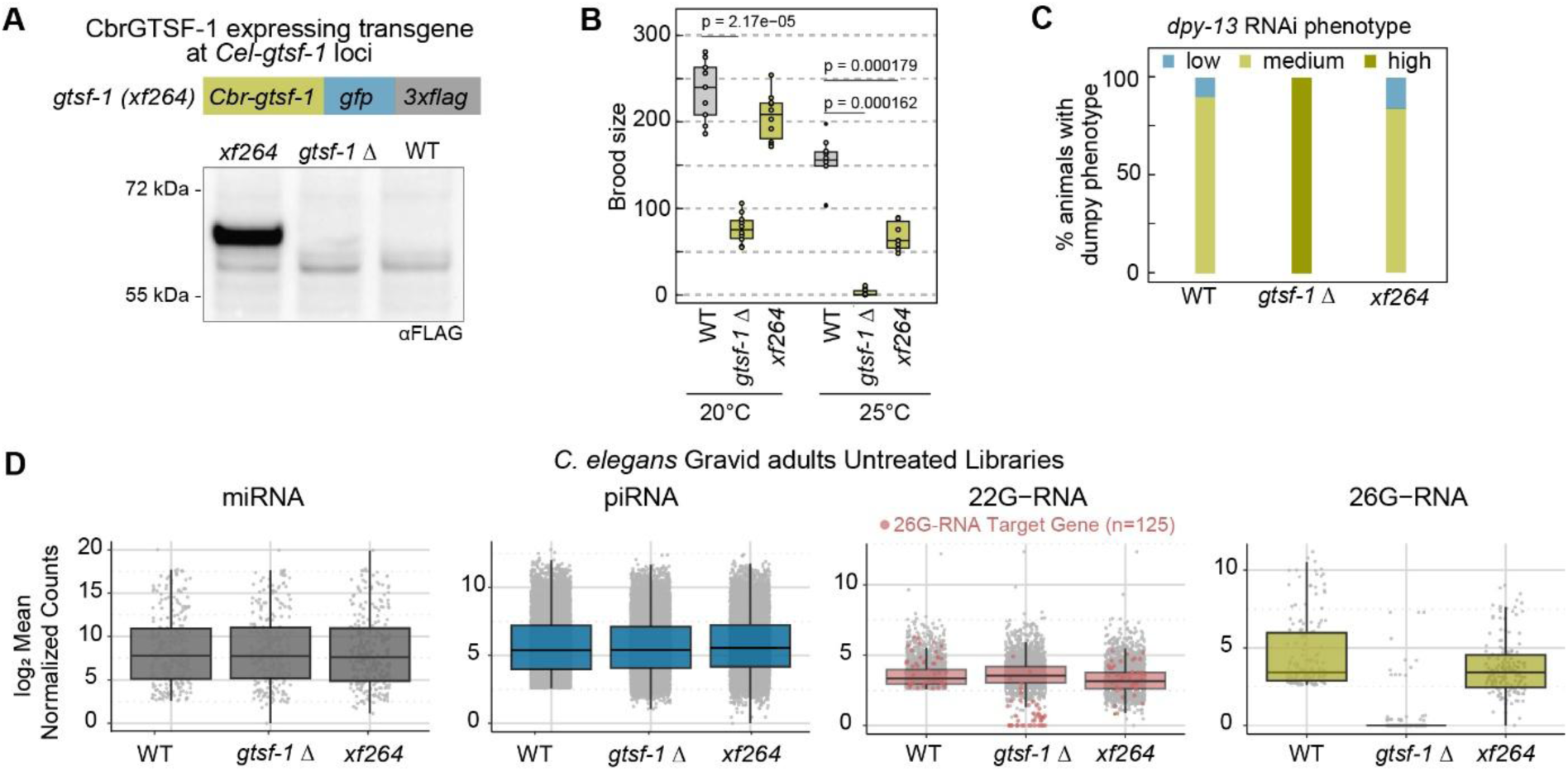
: Transgenic *C. briggsae gtsf-1* partially rescues *C. elegans gtsf-1* deletion (A) Top: Schematic of *Cel-gtsf-1(xf264)* allele with endogenous *gtsf-1* replaced by *C. briggsae gtsf-1* transgene upstream of *gfp* and *3×flag*. Bottom: Western blot from gravid adults validating transgene expression. (B) Brood size quantification (n = 15-25). P-values: Wilcoxon rank-sum test with BH correction. (C) RNAi responses to *dpy-13* scored by Dumpy phenotype penetrance and severity (low, medium, high) based on the visual ratio of body length to thickness. High severity: L2-like length with extreme thickening causing immobility. (D) Normalized counts per gene for small-RNA types. DESeq2-normalized counts averaged across three replicates. Colored 22G-RNA counts: 26G-RNA target genes identified in this library.

**Extended Figure 1.**
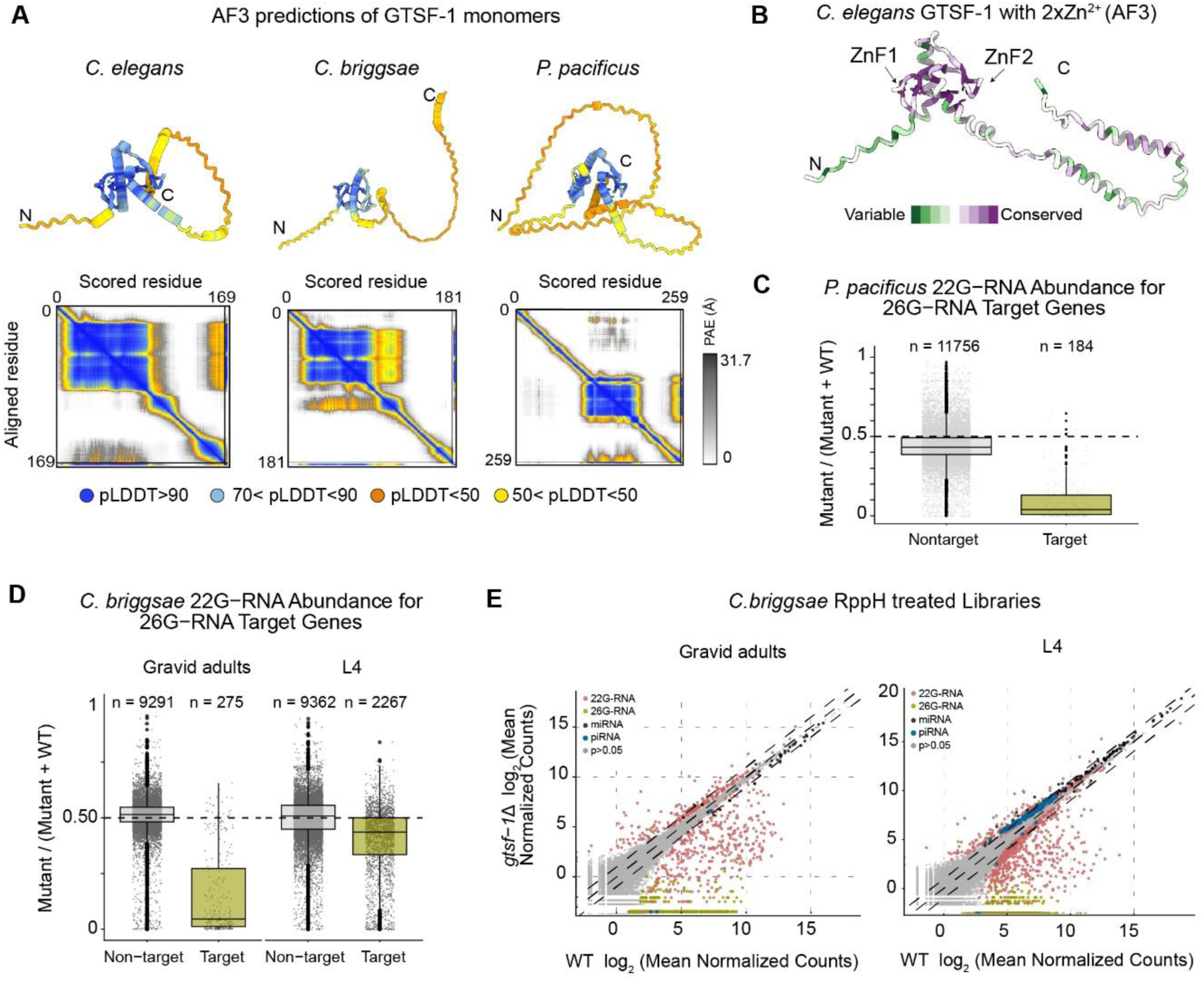
GTSF-1 structure prediction confidence and small-RNA sequencing in *gtsf-1* mutants (A) AlphaFold 3 monomer prediction of GTSF-1 colored by pLDDT score (0-100). ‘N’ and ‘C’ denote N and C-terminal residue. Bottom: Predicted aligned error (PAE) plots, regions colored by pLDDT score. (B) ConSurf conservation scores mapped onto *C. elegans* GTSF-1 AF3 structure using ClustalW MSA of 26 nematode species. Details in Table 2. (C-D) Abundance of normalized 22G-RNA counts per gene between wild-type and *gtsf-1* deletion samples. RppH-treated libraries were generated from (C) *P. pacificus* embryos or (D) *C. briggsae* gravid adults or L4 animals. Genes are classified as target of 26G-RNAs or non-targets. Dotted line: equal expression. n = number of genes per category. (E) Small-RNA counts per gene in RppH-treated libraries comparing wild-type and *gtsf-1* deletion animals in *C. briggsae* gravid adults and L4larvae. Colored points: p &lt; 0.05 (DESeq2, three replicates). Solid line: equal expression; dotted lines: ±2 fold change.

**Extended Figure 2.**
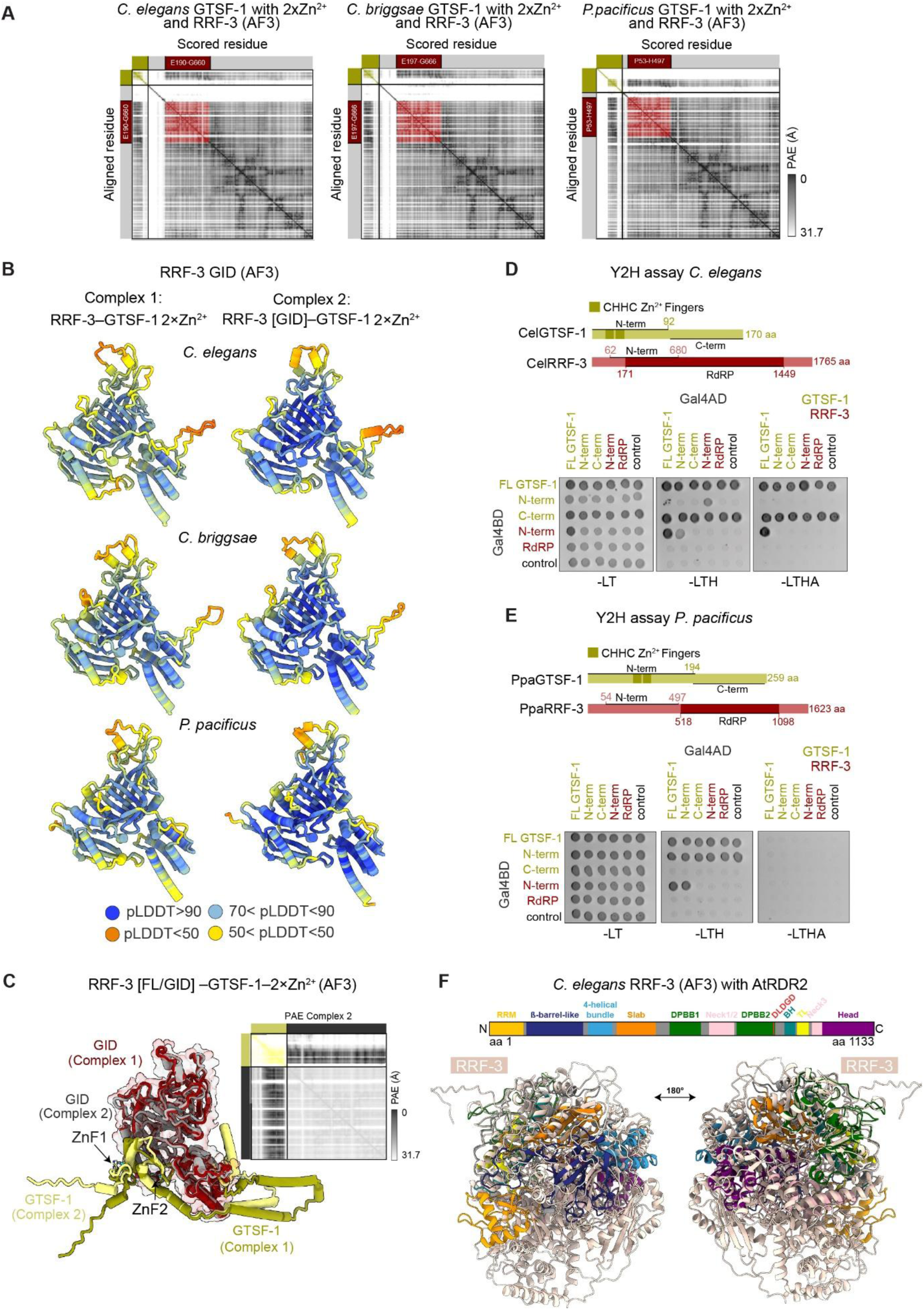
AlphaFold3 multimer models of nematode GTSF-1 and RRF-3 (A) PAE plots for GTSF-1-RRF-3-2×Zn²⁺ complexes. Red box indicates GID region and numbers indicate residues spanning the GID. Color scheme as Figure 4A. (B) Isolated RRF-3 [GID] structures (AF3) colored by pLDDT (0-100). Complex 1 = full-length RRF-3-GTSF-1-2×Zn²⁺, Complex 2 = RRF-3[GID] -GTSF-1-2×Zn²⁺. (C) *C. elegans* RRF-3 [GID] and GTSF-1 binding interface from complex 1 superimposed with complex 2, shows identical positioning of GTSF-1 N-terminal region. Right: PAE plot for Complex 1. (D, E) Yeast Two-Hybrid assays between GTSF-1 and RRF-3 regions under control (-LT: -Leu, -Trp), low stringency (-LTH: -Leu, -Trp, -His) and high stringency (-LTHA: -Leu, -Trp , -His, -Ade) selection. Control: Gal4 activation domain (AD) or DNA-binding domain (BD). (F) Top: Domain architecture of *Arabidopsis thaliana* RDR2from Figure 4E. Bottom: Superimposition of *C*. *elegans* RRF-3 on AtRDR2 with highlighted domains.

**Extended Figure 3.**
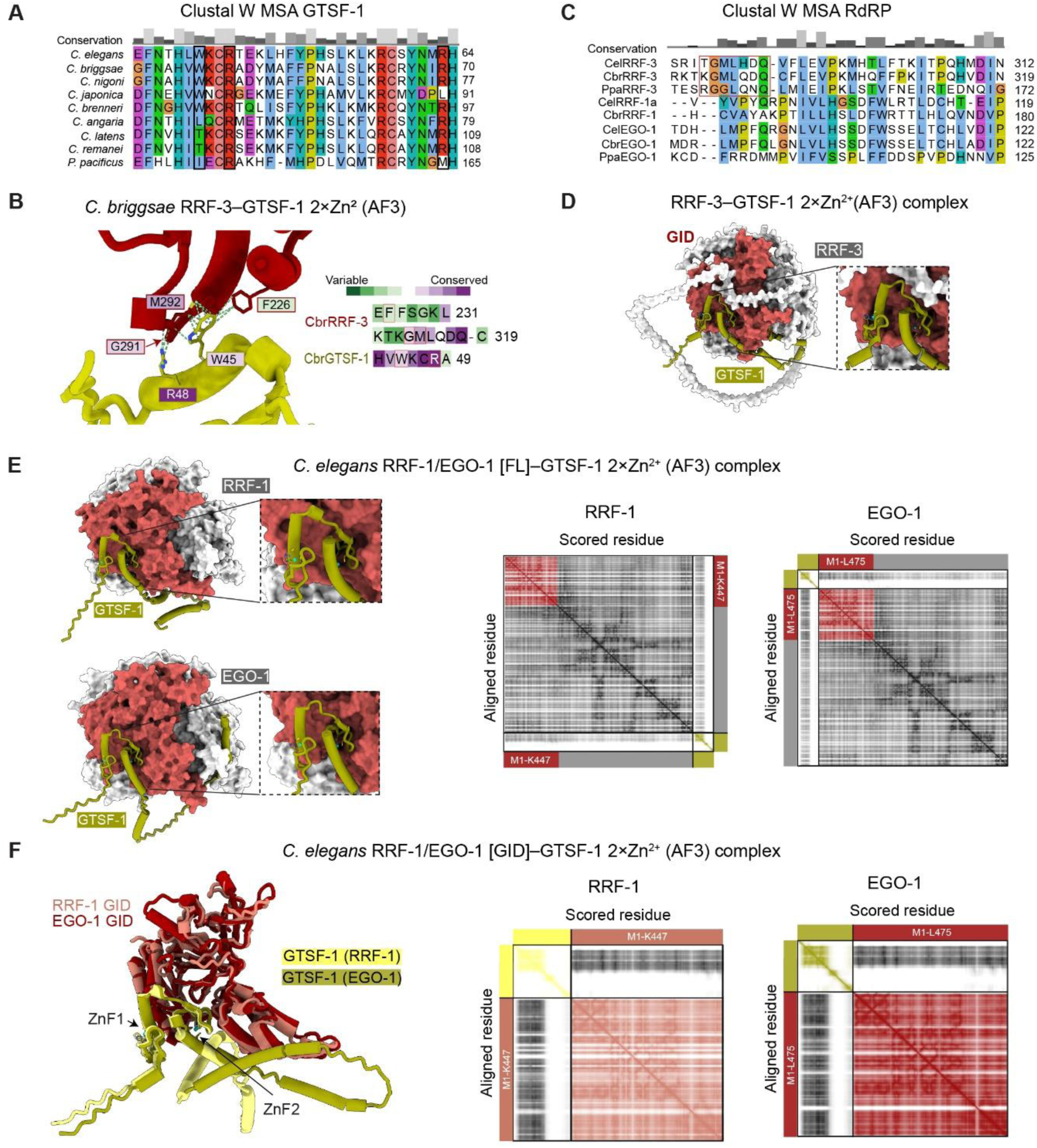
AF3 predicts a conserved interaction mode of GTSF-1 with RRF-1 and EGO-1 (A) Clustal Omega MSA of nematode GTSF-1 with tested residues from Figure 5A highlighted. (B) AF3 *C. briggsae* GTSF-1-RRF-3-2×Zn²⁺ interface with selected residues colored by ConSurf score (26 nematode MSA, as Extended Figure 1B). (C) Clustal Omega MSA of nematode RdRP proteins, highlighting conservation of residues in RRF-3 GID present in GTSF-1 binding interface. (D) AF3multimer prediction of *C. elegans* RRF-3 with GTSF-1 (two Zn²⁺ ions). The GID is highlighted in red. Inset: zinc finger docking of GTSF-1 onto RRF-3 GID surface. (E) Left: AF3 structures of *C. elegans* RRF-1 and EGO-1 showing GID interaction with GTSF-1 (two Zn²⁺ ions). Right: PAE plots for the respective models (F) Left: Superimposed AF3 structures of *C. elegans* RRF-1 and EGO-1 isolated GID showing interaction with GTSF-1 (two Zn²⁺ ions). Right: PAE plots for the respective models.

**Extended Figure 4.**
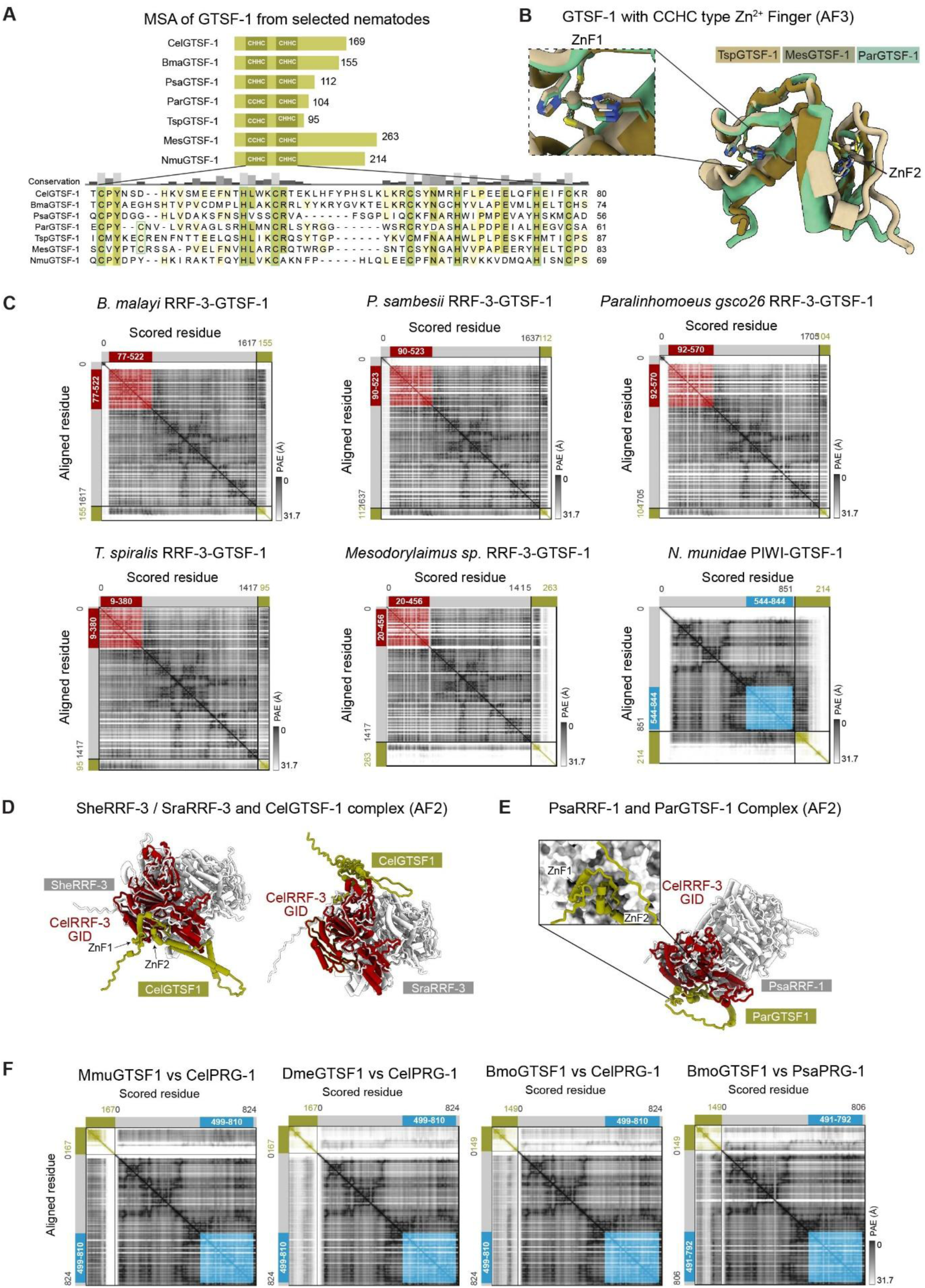
GTSF-1 structure and interaction mode with RRF-3 are conserved across the nematode phylum (B) MSA of GTSF-1 zinc finger regions from tested nematode species. Colored by percentage identity. Green boxes highlight CHHC residues. Species: Cel, C. elegans; Bma, *B. malayi*; Psa, *P. sambesi*; Par, *Paralinhomoeus gsco26*; Tsp, *T. spiralis*; Mes, *Mesodorylaimus sp*.; Nmu, *N. munidae*. (C) Superimposed AlphaFold3 monomer structures of *T. spiralis* (Tsp), *Mesodorylaimus sp*. (Mes), and *Paralinhomoeus gsco26* (Par) GTSF-1 with two Zn²⁺ ions, highlighting CCHC type zinc fingers. (D) PAE plots of AlphaFold2 structures from Figure 6C. Red boxes highlight GID region with positions of spanning residues. Similarly for *N. munidae*, blue boxes highlight PIWI domain with positions of spanning residues. (E) AF2 multimer of RRF-3-GTSF-1 from *S. hermaphroditum* (She) and *S. ratti* (Sra) superimposed with *C. elegans* RRF-3[GID]. Green outline in *S. ratti* RRF-3 indicates absent regions of GID. (F) AF2 multimer of *Paralinhomoeus gsco26* (Par) and GTSF-1 *P. sambesi* (Psa) RRF-1, superimposed with *C. elegans* RRF-3 GID. Inset: zinc finger docking of ParGTSF-1 onto PsaRRF-1 surface. (G) PAE plots of AlphaFold2 structures from Figure 6E.

